# Histone’s quality control by prokaryotic ClpP/ClpR regulates the eukaryotic mitotic cell cycle in malaria parasites

**DOI:** 10.64898/2026.05.13.718385

**Authors:** Sudipta Das

## Abstract

In *Plasmodium falciparum*, DNA replication and asynchronous nuclei formation precede cytokinesis during intraerythrocytic schizogony. Inhibition of fatty acids (FAs) import and impaired membrane biogenesis led to the arrest of mitosis through the inhibition of DNA replication and nuclei formation. On the iRBC surface, parasite ribosomal protein P2 (PfP2) complex mediated FAs import and membrane biogenesis, seemingly prior events before the commitment for DNA replication and nuclei formation. The inhibition of FAs import led to the degradation of histones by the evolutionarily conserved bacterial serine protease ClpP/ClpR in the parasite nucleus. Noncanonical arginine hyperphosphorylation by a novel arginine kinase in the nucleus was subjected for proteostasis and marks histones for degradation by ClpP/ClpR machinery. Inhibition of *de novo* FAs biosynthesis by an anti-cancer drug, Cerulenin and C75, in HEK293T and HCT116 carcinoma mammalian cells showed histone degradation. Lipid (L) induced histone proteostasis by ClpP/ClpR, seemingly an indispensable L-checkpoint before mitotic commitment.

## Introduction

*Plasmodium falciparum* (*P. falciparum*), the causative agent of human malaria, remains a huge health burden globally, mostly across developing countries^1^. During intraerythrocytic schizogonic cell division, *P. falciparum* shows repeated asynchronous nuclear division and nuclei formation, giving rise to 16–32 merozoites ^2, 3, 4, 5, 6, 7, 8, 9^. During asynchronous nuclear division, repeated DNA replication and mitosis (S/M Phase) precede cytokinesis and merozoite formation ^7, 9^. At the onset and during ongoing nuclei formation, import of fatty acids (FAs) and membrane biogenesis are critical steps for cell cycle progression ^10,11^. The inhibition of FAs import resulted into the cessation of DNA replication and nuclei formation at the onset of schizogony ^12^. In any eukaryotic cellular species, availability and organization of membrane precursors and bi-layer assembly provide a necessary confirmation for cells to go ahead and commit to mitotic division ^13,14,15,16,17^. Hence, the scarcity of any precursor molecules required for membrane assembly in principle may initiate a signalling cascade to cease the cell cycle before commitment or during ongoing mitotic progression ^17^. At the trophozoite stage, PfP2 complex interacts with the Skeletal Binding Protein 1 (SBP1) in the infected RBC (iRBC) cytosol and translocates to the iRBC surface for importing FAs, which is necessary for downstream synthesis of membrane components and bi-layer assembly ^12,18^. Inhibition of PfP2 complex by PfP2 specific monoclonal antibody E2G12 inhibited FAs import and resulted into the arrest of nuclear division at the onset of schizogony ^12^. To explore how the inhibition of FAs import led to the arrest of mitotic schizogony, we have characterized the signalling pathway using total transcriptomics, RNA sequencing, lipidomics, and proteomics approaches and discovered that the inhibition of FAs import led to the degradation of parasite histone proteins by an evolutionarily conserved prokaryotic protein quality control system, ClpP/ClpR, in the parasite nucleus. In bacteria, serine protease ClpP is conserved and has been identified to perform protein quality control ^19,20,21^. Evolutionarily, ClpP is conserved across cellular systems from prokaryotes to eukaryotic cells and found to be operational in cellular organelles, e.g., in mitochondria of mammalian cells for protein quality control ^22,23^. Unequivocally, FAs biosynthesis and membrane assembly are fundamental cellular processes for cell division, and all cell types require FAs for the synthesis of membrane components. When we checked the effect of the inhibition of *de novo* biosynthesis of FAs in cultured HEK293T and HCT116 carcinoma mammalian cells using FAs biosynthesis inhibitor Cerulenin and C75, similar to *P. falciparum*, astonishingly, we observed the degradation of histone H2A, H2B, H3, and H4, resulting in cell cycle arrest (Figure S1).

In *P. falciparum* 3D7 and NF54 strains, inhibition of FAs import and the scarcity of membrane components led to the hyperphosphorylation of arginine in all histones, H2A, H2B, H3, and H4. The hyperphosphorylation of arginine marks histones for degradation by nuclear localized ClpP/ClpR quality control machinery 19. Astonishingly, in the parasite, the ability of hyperphosphorylation of arginine residues in histones and recognizing hyperphosphorylated histones by ClpP/ClpR was transferable to mammalian histones, and consequentially degradation of mammalian histones H3 and H4 were also observed. We hypothesized that the degradation of parasite histones provides a necessary framework and signalling cascade to arrest DNA replication and nuclei formation through the inability of histone octamer formation and daughter nucleosome assembly at the onset of schizogony. The evolutionarily conserved ClpP/ClpR mediated histone’s quality control in the parasite nucleus provides a unique and novel mechanism of lipid-mediated mitotic checkpoint event and cell cycle regulation in human malaria parasites at the onset of schizogony. The ClpP/ClpR-mediated cell cycle checkpoint event in *P. falciparum* opens a new avenue for further investigation into whether this lipid-mediated mechanism of the mitotic checkpoint is conserved across all eukaryotic cell types at the onset and during ongoing cell cycle progression.

## Results

### Inhibition of FAs import led to the degradation of parasite histones

Our previous report in iScience ^12^ showed that the inhibition of FAs import in iRBCs resulted into the arrest of parasite cell cycle^12^. To unpack the reason for cell cycle arrest, we performed biochemical assays using cultured *P. falciparum* 3D7 and NF54 parasites. In Growth Inhibition Assay (GIA) (Figure. 1A), Anti-PfP2 monoclonal antibody (E2G12) ^12, 24, 25^ was used to block the import of serum fatty acids (FAs) via the blockage of FAs importing PfP2 protein complex on the iRBC surface ^12,24^. As a result, the parasite nuclear division was arrested at trophozoite stage and subsequently, when the parasites were rescued from the E2G12 treatment, the nuclear division proceeded towards schizogony (Figure. 1B). The E2G12 mediated inhibition of FAs import and subsequent effect as cell cycle arrest at trophozoite stage was quantitatively corroborated with known cell cycle arrester molecules (Figure. 1C). Interestingly, after the rescue from E2G12 treatment, arrested parasites proceeded towards schizogony (Figure. 1B & 1C). In Histone 3 (H3), acetylation of the 56^th^ Lysine (^56^K) residue (H3^56^K-ac) regulates the DNA replication, enhances transcription, and regulates DNA damage response in yeast ^26,27,28,29^. We wanted to check the expression level of H3 and H4 and the status of posttranslational modification (PTMs) of H3-^56^K residue as a consequence of FAs import inhibition. To our surprise, in the FAs import inhibited parasites, we observed a significant reduction in the level of parasite histone H3 and H4 (Figure. 1D). Interestingly, at the same trophozoite stage of parasite life cycle, other cell cycle arrester, e.g., Taxol, Aphidicolin and Orlistat did not show decreased level of H3 and H4 (Figure. 1D). In the E2G12 rescued parasites, H2A, H2B, H3 and H4 level came back to a significant level suggested the reason for the progression of nuclear division towards schizogony (Figure. 1D). Inhibition of FAs import and the treatment with Orlistat, both resulted into the upregulation of Triacylglycerol (TAG) in the parasite ^12^. Hence, merely enhanced TAG level did not appear to be the reason for histone degradation. In the life cycle of *P. falciparum* 3D7, time point analysis under GIA condition with E2G12, Post Antibody Incubation (PAI) 12h onwards (PMI~22-24h), FAs import inhibition through PfP2 blockage by E2G12 resulted in H3 and H4 degradation (Figure. 1E & 1G). In the parasite during blood stage infection, there is no evidence that suggests a *de novo* FAs biosynthesis, instead parasite is dependent on salvage from host cells and import from host serum^30, 31, 32, 34^. Parasite culture medium deprived of Palmitic acid or Oleic acid resulted into cell cycle arrest at trophozoite stage ^33, 34, 35, 36^. Imported FAs from extracellular sources assimilate into the parasite membrane during membrane lipid synthesis and bilayer assembly ^35, 36^. At PMI~24-26h, DNA replication and nuclear division begin; hence, the inhibition of FAs import presumably compromised membrane biogenesis, which was required for daughter nuclei formation. On the contrary, in *P. falciparum* NF54 life cycle, we observed an interesting opposite result. Under GIA condition with E2G12, we observed H3 and H4 degradation at PAI~12h (PMI~24h) and 24h (PMI’34h), but in the subsequent time points, at PAI 30h and 36h, the level of H3 and H4 came back even in the presence of E2G12, contrary to *P. falciparum* 3D7 parasites (Figure. 1F & 1H). This observation raised a question about how the H3 and H4 levels came back in the presence of E2G12 and whether it is somehow linked with the concomitant trigger for *de novo* biosynthesis of FAs in the apicoplast of *P. falciparum* NF54 parasites. In *P. falciparum* 3D7 transgenic parasites (Pf3D7P2-3xHA-glmS)^12^, the downregulation of PfP2 protein in the presence of 3mM Glucosamine (GlcN), resulted into the degradation of H3 and H4 (Figure. 1I). The downregulation of PfP2 protein caused the inhibition of the formation of FAs importing complex on the iRBC surface which resulted into FAs import inhibition, similar to blockage through E2G12 antibody ^12^. In *P. falciparum* NF54 transgenic parasites (PfNF54 P2-3xHA-glmS-Ribozyme), when FAs import was inhibited through the genetic downregulation of P2 protein using 3 mM GlcN, we also observed degradation of histone H3 and H4 at the trophozoite stage (PMI~24h) (Figure. 1J, Figure. S2). Both the results corroborated with the observations that the upstream downregulation of PfP2 protein plausibly led to the malformation of FAs importing complex on the iRBC surface, which resulted into the inhibition of FAs import and subsequently compromised membrane biogenesis and consequentially the degradation of H2A, H2B, H3, and H4. These entire results posed a legitimate question whether it was actually histone degradation or transcriptional downregulation or both as a result of FAs import inhibition. If it was a degradation then what caused it, and what was the mechanism?

**Figure 1.**
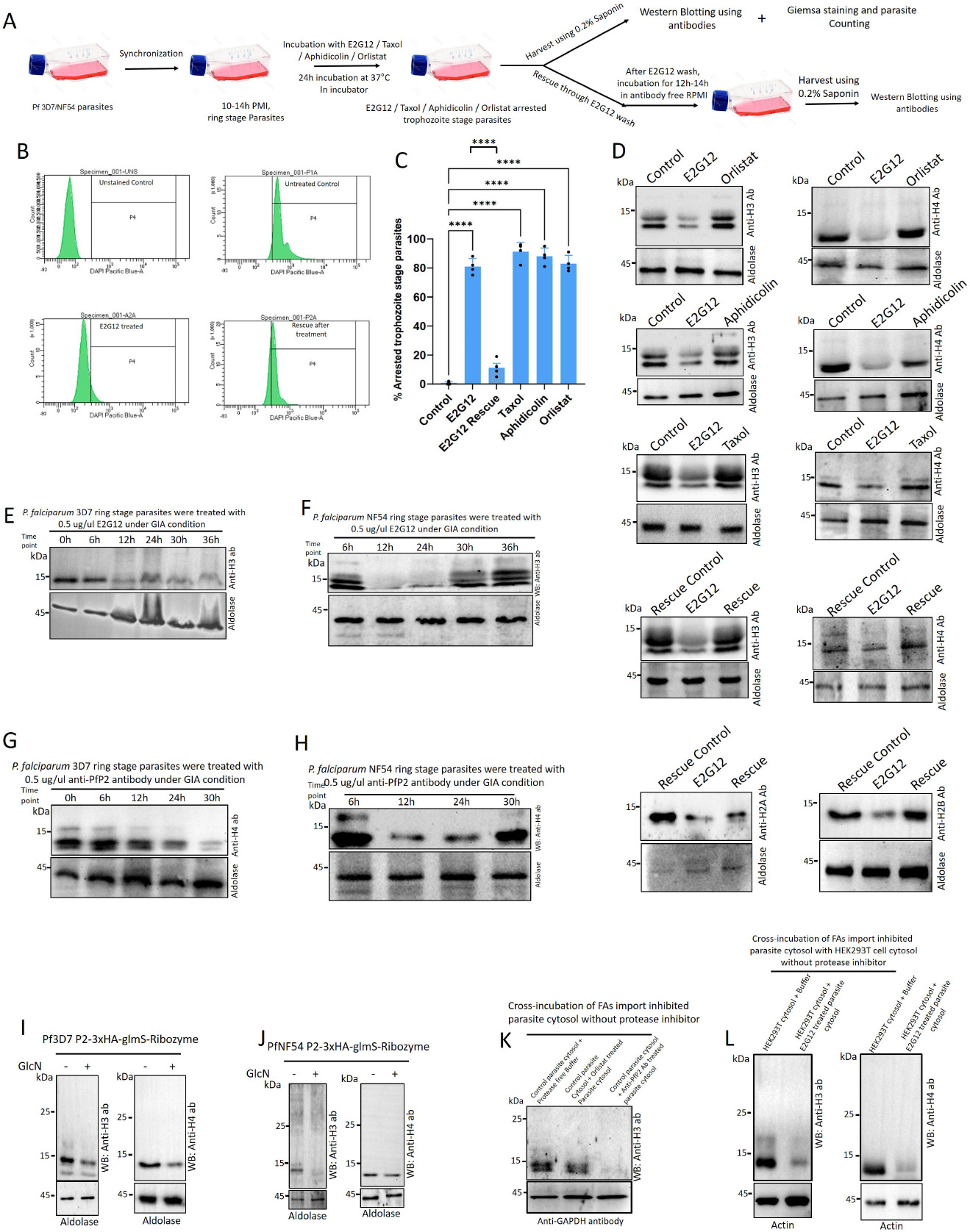
Inhibition of FAs import led to arrest of parasite nuclear division due to Histone degradation. **(A)** Schematic showing the strategy of growth inhibition assay (GIA), rescue and harvest of *P. falciparum* 3D7 and NF54 parasites using anti-PfP2 specific monoclonal antibody E2G12, Taxol, Aphidicolin and Orlistat. **(B)** Flow cytometric analysis of *P. falciparum* 3D7 nuclei formation using nuclear stain DAPI under control (untreated), E2G12 treated and E2G12 free rescued condition. Under each condition, 0.5 million iRBCs were counted for DAPI positive cells and plotted. After treatment, iRBCs were harvested at PMI~ 34h for DAPI staining and counting. **(C)** Quantification of cell cycle arrested *P. falciparum* 3D7 parasites at trophozoite stage under control (untreated), E2G12 treated, E2G12 free rescued, Taxol, Aphidicolin and Orlistat treated condition. Biological replicates N=4. Statistical P values were denoted as *. P<0.05 as *, P<0.01 as **, P<0.001 as *** and P<0.0001 as ****. **(D)** Immunoblot (Western blot) analysis of *P. falciparum* 3D7 parasites at trophozoite stage under control (untreated), E2G12 treated, E2G12 free rescued, Taxol, Aphidicolin and Orlistat treated condition. Under different treated conditions, parasite histone 2A (H2A), histone 2B (H2B), histone 3 (H3) and histone 4 (H4) were probed and parasite Aldolase was used as a loading control. **(E & G)** *P. falciparum* 3D7 parasites at PMI ~12-14h were treated with E2G12 (700 ng/µl) under GIA condition at different time points up to 36h post E2G12 addition in the culture media. Post addition of E2G12 in the culture media, after every 6h at different time points parasites were harvested and immunoblot was done using histone 3 (H3) and histone 4 (H4) antibody. Parasite Aldolase was used as a loading control. (F & H) *P. falciparum* NF54 parasites at PMI ~12-14h were treated with E2G12 (700 ng/µl) under GIA condition at different time points up to 36h post E2G12 addition in the culture media. Post addition of E2G12 in the culture media, after every 6h at different time points parasites were harvested and immunoblot was done using histone 3 (H3) and histone 4 (H4) antibody. Parasite Aldolase was used as a loading control. (I) Synchronized *P. falciparum* 3D7 PfP2-HA glmS-Ribozyme (Das et al., 2024)^12^ transgenic parasites were treated with 3 mM glucosamine (GlcN) for PfP2 downregulation. In the same parasite line after PfP2 downregulation, immunoblot was done to check the level of H3 and H4 using H3 and H4 antibody. As a loading control, parasite Aldolase was use. (J) Synchronized *P. falciparum* NF54 PfP2-HA glmS-Ribozyme transgenic parasites were treated with 3 mM glucosamine (GlcN) for PfP2 downregulation. In the same parasite line after PfP2 downregulation, immunoblot was done to check the level of H3 and H4 using H3 and H4 antibody. As a loading control, parasite Aldolase was use. (K) Doubly synchronized *P. falciparum* 3D7 parasites were treated with control (untreated) and 700 ng/µl E2G12 and Orlistat (20 µM) for 24h. The cytosol of treated parasites was harvested under protease-free condition. The cytosol was cross-incubated where control (untreated) parasite cytosol was incubated with Orlistat treated parasite cytosol, control (untreated) parasite cytosol was incubated with E2G12 treated parasite cytosol and as a control, the control (untreated) parasite cytosol was incubated with proteas free buffer. The cross-incubation of parasite cytosol was done at 37°C for 6h. Total 2 µg of total cytosol protein was incubated at 1:1 ratio. Incubated cytosols were probed in immunoblot with anti-Histone 3 (H3) antibody. GAPDH was used as a loading control. (L) The cytosol of HEK293T cultured mammalian cells were cross-incubated with E2G12 treated *P. falciparum* 3D7 parasite cytosol under protease free condition at 37°C for 6h. Total 2 µg of total cytosol protein was incubated at 1:1 ratio. As a control, HEK293T cell cytosol was treated with protease free buffer. Incubated cytosols were probed in immunoblot with anti-histone 3 (H3) antibody. Actin was used as a loading control.

Presumably, degradation of H2A, H2B, H3, and H4 was achieved either through histone poly-Ubiquitination and proteasome mediated or resulted due to the activation of a Protease. In immunoblot data using H2A, H2B, H3, and H4 specific antibodies, we did not see higher ubiquitinated histone bands, neither in parasites after E2G12 mediated arrest nor in the mammalian cells after Cerulenin and C75 treatment (Figure. 1I, 1J & S1A). It suggested that in the parasites, enhanced TAG level did not cause ubiquitination of histones. Now we were left with the possibility of a protease. To check the authenticity of a protease-mediated histone degradation, we did not rely only on the observation of histone degradation in the GIA assay; instead, we checked the possibility of the presence of a protease in the E2G12 treated parasite cytosol by cross-incubating control parasite cytosol with E2G12 treated parasite cytosol. To our surprize, H3 in control parasite cytosol was degraded in the presence of E2G12 treated parasite cytosol (Figure. 1K). It did indicate that when FAs import was blocked by inhibiting PfP2 complex on the iRBC surface using E2G12, a histone specific protease was getting upregulated and as a result, H3 in the E2G12 arrested parasites was degraded. In addition, we also wanted to check whether this ability of histone degradation was transferable to other cell types and this unknown protease has the ability to degrade mammalian histones as well. To ascertain that, the cytosol of mammalian cell HEK293T was cross-incubated with E2G12 arrested parasite cytosol followed by immunoblotting. To our surprize, H3 and H4 of HEK293T cells were significantly diminished as compared to untreated samples (Figure. 1L). This seemed to suggest that due to the lack of FAs import through the blockage of PfP2 protein complex on the iRBC surface, a histone specific protease acquired the general ability to degrade histones. Degradation of mammalian histones by Plasmodium protease has opened a new avenue to look at the cross-talk mechanism between cell cycle regulation and membrane biogenesis through FAs import.

Thereafter, we wondered about the possibility of histone degradation in cultured mammalian cells, HEK 293T and HCT 116, when *de novo* biosynthesis of FAs was inhibited by the anti-cancer drug Cerulenin and C75. Astonishingly, all histones, H2A, H2B, and H3 and H4 were degraded when *de novo* FAs biosynthesis was disrupted (Figure. S1A). Further, we wanted to check whether DNA replication was halted or still continuing when *de novo* biosynthesis of FAs was inhibited by Cerulenin and C75. Flow cytometry analysis seemed to suggest a significantly reduced number of cells in treated samples, but surprisingly, Mean Florescence Intensity (MFI) of nuclear stain DAPI between untreated (control) and Cerulenin and C75 treated cells did not change significantly (Figure. S1B). This paradoxical result suggested that probably, DNA replication continued but FAs deprivation led to the inhibition of membrane biogenesis, and hence, cytokinesis was affected and consequentially cell division did not progress.

### Inhibition of FAs import triggered gametocytogenesis in *P. falciparum* NF54

Serum lyso-phosphatidylcholine (LysoPC) and the regulation of its import into the parasite during blood stage infection regulate sexual stage gametocytogenesis in *P. falciparum* ^37, 10.^ Since FAs are the precursor molecules for the biogenesis of PC through the Kennedy Pathway ^38, 39^, we wondered about the biochemical and morphological transformations of *P. falciparum* NF54 parasites when FAs import was blocked through the inhibition of the PfP2 complex on the iRBC surface using anti-PfP2 mAb E2G12. To ascertain this hypothesis, we performed time point experiments with *P. falciparum* NF54 parasites to assess morphological transformations and gametocyte formation. Upon inhibition of FAs import, PAI 34h onwards, gametocytes were observed, and at later stages, PAI 40h, more mature male and female gametocytes were significantly higher (Figure. 2A & 2D). Whereas, preimmune and untreated parasites did not show gametocyte production but Orlistat treatment appeared to induce 10-15% somewhat elongated parasites (Figure. 2A & 2D). When the P2 protein in *P. falciparum* NF54 parasites was downregulated using 3 mM GlcN, we also observed a significant morphological switch and gametocyte induction (Figure. S2E). We questioned as to how much biochemistry took place, which led to this remarkable gametocyte transformation. Was it the case that at later time points, somehow the level of histone H3 and H4 was getting rescued even under continuous inhibition of FAs import through PfP2 complex on the iRBC surface by the presence of E2G12 in the culture media (Figure. 1F & 1H). This may only be possible if *de novo* biosynthesis of FAs was activated through gene regulation to circumvent the FAs need to restore membrane biogenesis and rescue the level of H3 and H4 to proceed with DNA replication and transcription regulation. Hence, as a result, it might signal for morphological transformation of the parasites and end up forming gametocytes. To ascertain our hypothesis, we approached the problem through total lipid analysis (lipidomics) and mRNA sequencing and total transcriptomics. LC-MS lipid profile of *P. falciparum* NF54 parasites after E2G12 treatment at PAI 40h showed few unique lipid species in the treated parasites e.g., sterol lipid (m/z 371.1823/372.1883 Da), LCB-Sphingolipids (m/z 337.1647) (Figure. 2B & 2C) (Excel sheet. S1 & S3). The biosynthesis of LCBs involves complex pathways starting from serine and palmitoyl-CoA^40, 41^. LCBs are combined with a fatty acid and, in many cases, another group (like a phosphate or sugar) to form a variety of sphingolipids, such as ceramide. The synthesis of LCBs begins with serine and palmitoyl-CoA, catalysed by the enzyme palmitoyltransferase ^40, 41^. These observed unique lipids suggested a transcriptional trigger which was necessary to initiate a pathway to synthesize lipid species for gametocyte production.

**Figure 2.**
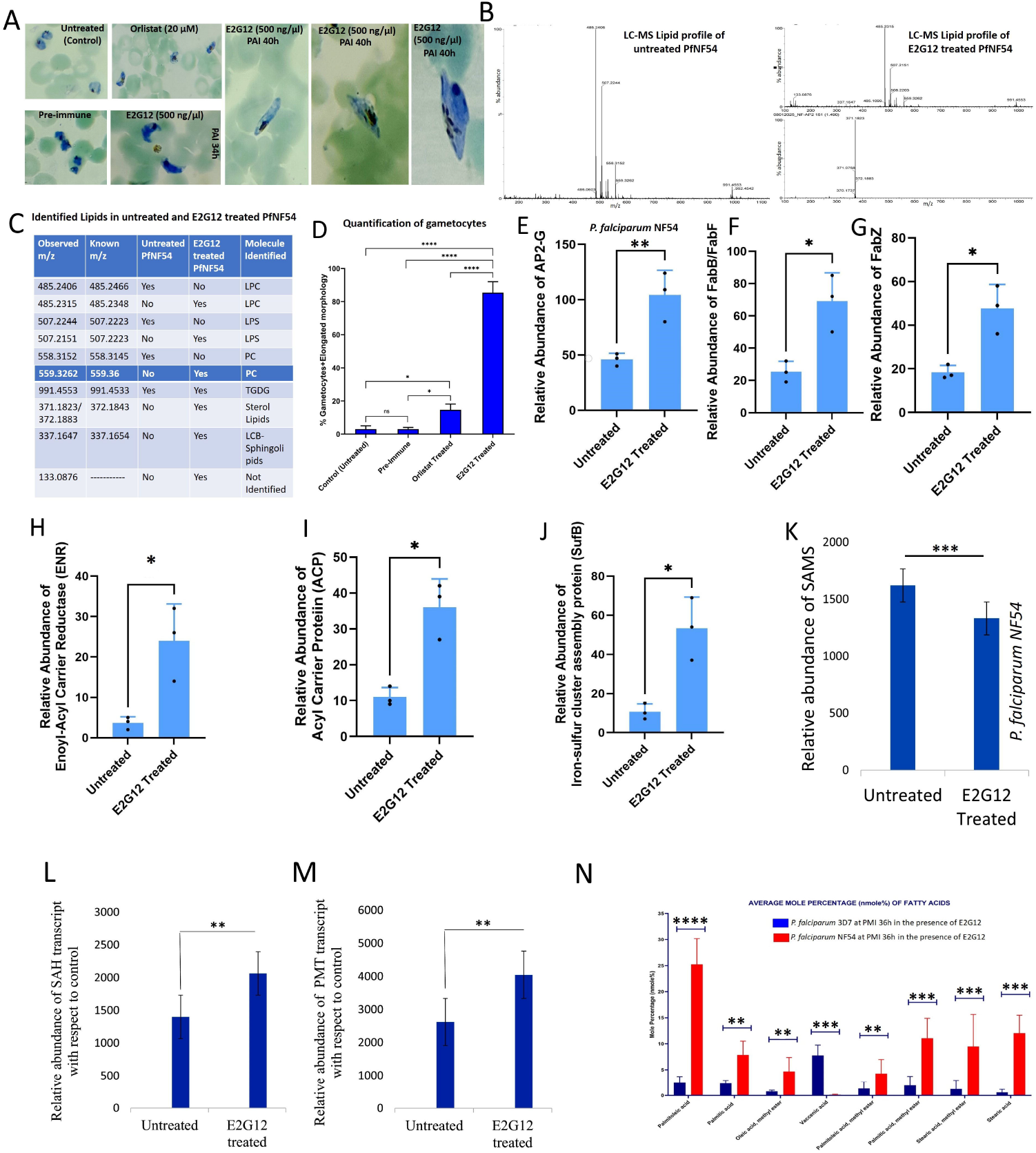
FAs import inhibition triggered *de novo* FAs biosynthesis in the apicoplast and induced gametocytogenesis in *P. falciparum* NF54. **(A)** Double synchronized *P. falciparum* NF54 parasites were treated with Control (untreated), pre-immune rabbit sera, 20 µM Orlistat and E2G12 (700 ng/µl) at 37°C for Post Antibody Incubation (PAI) 34h and 40h. Parasites were stained with Giemsa. Shape of the parasites were examined by microscopy and quantified. **(B)** LC-MS profile of Lipids of *P. falciparum* NF54 parasites at trophozoite stage under control (untreated) and E2G12 (700 ng/µl) treated PAI 40h at 37°C. X axis showing m/z and Y axis depicts % abundance. **(C)** Identified Lipid species in *P. falciparum* NF54 parasites at trophozoite stage under control(untreated) and E2G12 treated condition. **(D)** Quantification of parasite morphology under control (untreated), pre-immune sera treated, 20 µM Orlistat treated and E2G12 (700 ng/µl) treated at 37°C. X axis showing treatment condition and Y axis showing % gametocytes+ elongated morphology. **(E-J)** Relative transcripts abundance of AP2-G, FabB/FabF, Enoyl-acyl carrier reductase (ENR), Acyl-carrier protein (ACP) and Iron-sulphur cluster assembly protein (SufB) in *P. falciparum* NF54 parasites at trophozoite stage under control (untreated) and E2G12 (700 ng/µl) treated condition at 37°C PAI 34h. **(K-M)** Relative abundance of transcripts of S-adenosylmethionine synthetase (SAMS), S-adenosyl-L-homocysteine (SAH) and Phosphoethanolamine methyltransferase (PMT) in untreated Vs E2G12 treated (arrested at trophozoite stage) parasites (N) Mole percentage (nmole %) of Fatty Acid (FAs) abundance in *P. falciparum* 3D7 and NF54 parasites under E2G12 (700 ng/µl) treated condition at 37°C PMI 36h. X axis showing types of FAs and Y axis is mole% (nanomole%). N=3. Statistical P values were denoted as P<0.05 as *, P<0.01 as **, P<0.001 as *** and P<0.0001 as ****.

Gametocyte induction is regulated by a master transcription factor AP2-G ^42, 43^. When *P. falciparum* NF54 parasites were treated with E2G12, causing inhibition of FAs import, the relative abundance of AP2-G transcripts (PF3D7_1222600) was significantly upregulated (Figure. 2E, Excel sheet. S2). In addition, transcripts related to *de novo* FAs biosynthesis in the apicoplast genome, e.g., FabB/FabF, Enoyl acyl-carrier Protein (ENR), Acyl-carrier Protein (ACP), and Iron sulphur cluster assembly protein (SufB), were also significantly upregulated (Figure. 2E-2J, Excel sheet. S2). Interestingly after Orlistat treatment, neither AP2-G nor *de novo* FAs biosynthesis-related genes in the apicoplast were upregulated (Excel sheet. S3). This suggested that merely enhanced Triacylglycerol (TAG) was not enough to trigger gametocytogenesis. All together, these suggested that the inhibition of FAs import led to dual transformations, morphological (shape change through gametocyte formation) and molecular (circumvent to mitigate the availability of FAs through *de novo* biosynthesis). In *P. falciparum* NF54, Phospho-ethanolamine methyltransferase (PMT) activity regulates the intracellular level of S-adenosylmethionine (SAM) and S-adenocyl homocysteine (SAH) ^44^. Reduction in intracellular SAM induced a 2-3-fold increase in sexual commitment for gametocyte formation ^44^. In our study, when FAs import was inhibited, we observed a significant upregulation of SAH and PMT and a reduction in SAM level, which corroborated with the previous study ^44^ and further confirmed that FAs import inhibition triggered sexual commitment and induced gametocytogenesis (Figure. 2K, 2L & 2M, Excel sheet. S2). Under GIA condition, in the presence of E2G12, PMI~36h, mol% of FAs and derivatives of different types of FAs, showed a significantly higher abundance in *P. falciparum* NF54 parasites as compared to *P. falciparum* 3D7, which indicated the initiation of *de novo* biosynthesis of FAs, the necessity and the importance of FAs availability and subsequent membrane biogenesis for gametocyte transformation (Figure. 2N).

Inhibition of FAs import led to the morphological transformation in *P. falciparum* NF54, which provided an indication of membrane composition shift for membrane assembly in male and female gametocytes (Figure. 2A). We hypothesized that the inhibition of FAs import changed lipid homeostasis through the synthesis of unique lipid molecules, which may played critical role in the morphological transformation and gametocyte specific membrane biogenesis. Synchronized *P. falciparum* NF54 parasites at PMI 12-14h were untreated or treated with pre-immune sera (700 ng/ul) and E2G12 (700 ng/ul) for 24h. Parasites were harvested, and total lipid was extracted for LC-MS analysis. In untreated and or pre-immune treated parasites, identified lipids were broadly classified as (1) low fatty acids conjugates (2) Glycero-lipids (3) Glycerophospholipids (4) Ceramides, whereas in the E2G12 treated parasites identified lipids were classifies as (1) relatively low fatty acids conjugates (2) Glycerophospholipids and (3) a unique peak (m/z 371.182) which was significantly abundant and it was identified as sterol lipids (Figure. 2B, Excel sheet. S4). Further, to understand the gradual changes in lipid homeostasis by E2G12 dose dependent incubation in synchronized *P. falciparum* NF54 parasites and total lipid identification by LC-MS (Excel sheet. S4), synchronized parasites were treated with various concentrations of E2G12. As compared to control, at E2G12 concentration 50 ng/ul and 100 ng/ul, we did not see any deviation in identified lipid species except one extra peak (m/z 122.153) (Excel sheet. S4). At E2G12 concentration 300 ng/ul, a drastic change in the lipid peaks was observed, and the identified number of LC-MS peaks was significantly enhanced, showing several new lipid molecules (Excel sheet. S4). At E2G12 concentration 300 ng/ul, identified lipid molecules were broadly classified as (1) a significant induction of fatty acids conjugates (2) sudden induction in Phosphosphingolipids (3) TAG of incremental difference in carbon length and unsaturation (4) Sterol lipids (5) Glycerophospholipids (6) Glycero-lipids. Very surprisingly, at E2G12 concentration 700 ng/ul and 1000 ng/ul, identified lipids were similar to control and 50 ng/ul and 100 ng/ul of E2G12, where identified lipid classes were (1) Glycerophospholipids (2) ceramides (3) low fatty acids conjugates (4) glycerol-lipids (Excel sheet. S4). This data suggested that at E2G12 concentration 300 ng/ul, *de novo* biosynthesis of FAs was switched on due to the sufficient blockage of extracellular FAs import by E2G12. Additionally, signalling molecule sphingolipids and significant enhancement of sterol lipids seemed to indicate the initiation of a biochemical cascade in the parasite, which was instrumental in alleviating the problem of FAs import inhibition from extracellular sources and additional need of lipids to sustain parasite morphogenesis through gametocyte specific membrane biosynthesis. Surprisingly, once this is achieved, the number of lipid species at E2G12 concentration 500 ng/ul and 1000 ng/ul was reduced to four classes, which suggested that once the transformation is achieved through the changes in lipid profiles, it was maintained thereafter. So, E2G12 concentration 300 ng/ul, and the blockage of FAs import at that concentration was critical in triggering a change in the lipid homeostasis to support parasite morphogenesis and gametocyte formation.

### Catch me if you can: RNA sequencing and total transcriptomics to identify histone degrading protease in FAs import inhibited parasites

Inhibition of FAs import and compromised membrane biogenesis led to the cessation of DNA replication and arrest of parasite cell cycle^12^. We wondered how the DNA replication was halted when FAs import was inhibited and consequentially TAG catabolism and membrane biogenesis were impaired? To pursue this daunting task and to identify a possible protease and the mechanism of H2A, H2B, H3, and H4 degradation, FAs import-inhibited *P. falciparum* 3D7 and NF54 parasites were subjected to total transcriptomic analysis. Synchronized *P. falciparum* 3D7 and NF54 parasites were treated with E2G12 for 24h, and as per the strategy (Figure. 3A), the harvested parasites were subjected to RNA isolation and RNA sequencing through total transcriptomics. On the other hand, E2G12 treated parasites were also rescued from the treatment and harvested parasites were also subjected to RNA sequencing and transcriptomic analysis (Figure. 3A). In FAs import-inhibited *P. falciparum* 3D7 parasites, the apicoplast organelle was severely affected. The heatmap of mRNA transcripts suggested that an apicoplast serine protease ClpM (a component of ClpP/ClpR protease machinery) was significantly upregulated by log2 4-5-fold in its expression level (Fig. 3B & 3C). In Addition, iron sulphur cluster assembly protein (SufB), Enoyl acyl-carrier reductase (ENR) in the apicoplast were also upregulated by log2 3-5-fold, whereas acyl-CoA binding protein was downregulated by log2 3-4-fold (Figure. 3C, Excel sheet. S5). Additionally, upregulation of 4 genes was also observed, namely, a conserved protein, PF3D7_1141100, an apoplast ribosomal Protein L2, API01500, a secreted ookinete protein, putative (PSOP2), PF3D7_1367800, and reticulocyte binding protein homologue 4 (RH4), PF3D7_0424200. These four genes were upregulated by log2 3-fold. More interestingly, two genes which were consistently upregulated were Dynein heavy chain, PF3D7_1122900 by log2 4-5-fold and cysteine repeat modular protein 4 (CRMP4), PF3D7_1475400 by more than log2 5-fold (Figure. 3B & 3C). It was not clear why in FAs import inhibited parasites, Dynein and CRMP4 genes were upregulated, and what their role (s) are in this condition. The histones transcripts H2A, H2B, H3, and H4 all were downregulated by log2 1-2-fold (Figure. 3C & 3D, Excel sheet. S5). In the FAs import inhibited *P. falciparum* 3D7 parasites, the overall upregulated, downregulated, and unchanged transcripts were measured in log2 fold change and represented in the Volcano plot and scatter plot (Figure. 3D).

**Figure 3.**
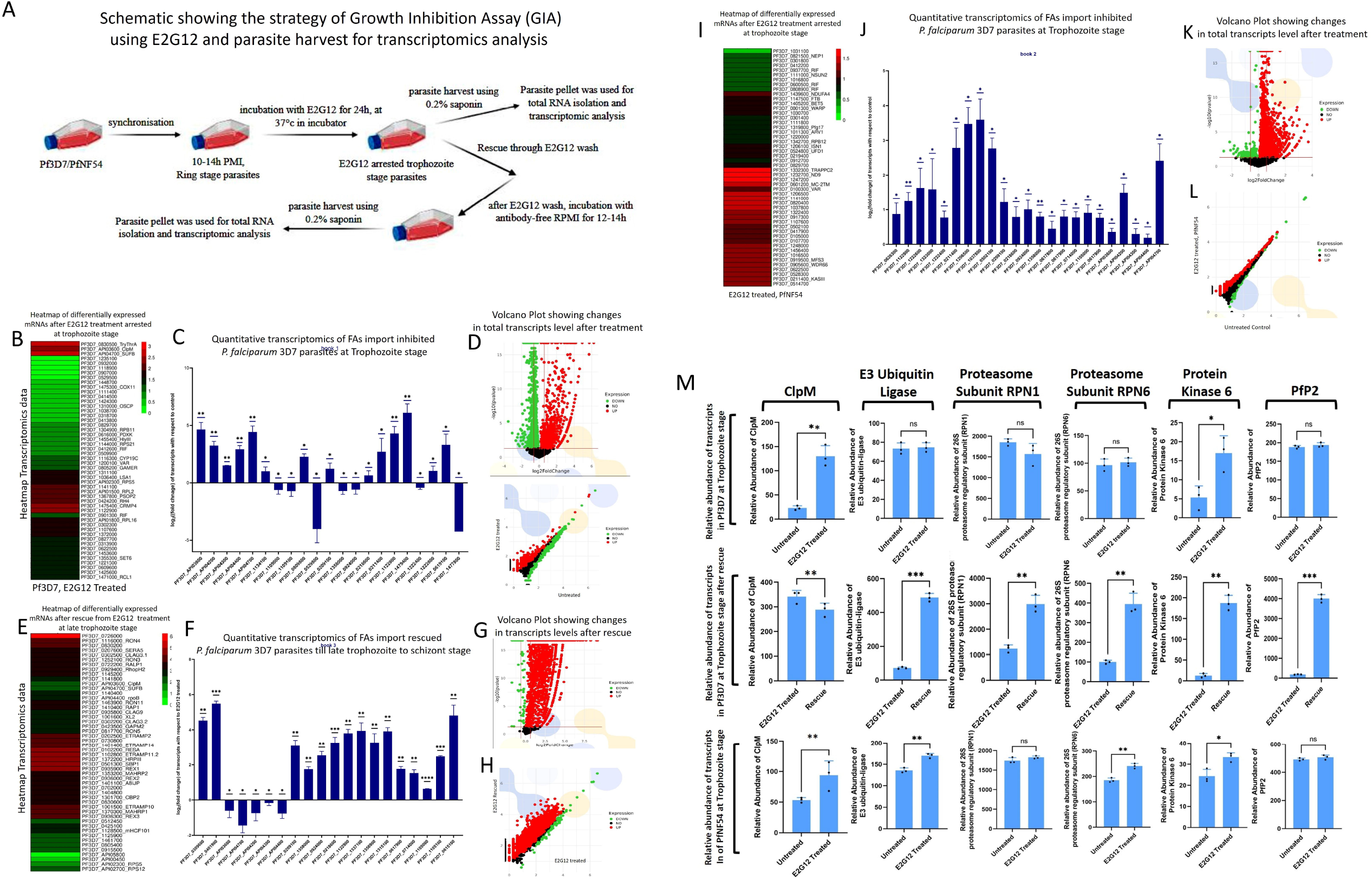
RNA sequencing and total transcriptomics to identify histone degrading protease and cascading events. **(A)** Schematic showing the strategy of Growth Inhibition Assay (GIA) of *P. falciparum* 3D7 trophozoite stage parasites using E2G12 antibody and parasite harvest for subsequent transcriptomics analysis. **(B)** Heatmap of 50 most differentially expressed *P. falciparum* 3D7 mRNAs at trophozoite stage after E2G12 treatment. Gene IDs were identified in plasmodb.org data base. The scale of the mRNA expression level was from 0 to 3, where 0 was comparable to control (untreated) parasites and 3 depicts the highest expression of the transcript. **(C)** Quantitative analysis of *P. falciparum* 3D7 mRNA transcripts by log2 fold change after E2G12 treatment at trophozoite stage. The upregulated and downregulated transcripts after E2G12 treatment were depicted by gene ID from plasmodb.org one a scale of log2 fold change from −5 to +10. mRNA transcripts were analysed on biological replicates N=3. **(D)** Volcano plot showing the changes in total mRNA transcripts after the treatment with E2G12. Each dot signifies a mRNA transcript. Green depicts downregulation of mRNA level; red depicts upregulation of mRNA and black depicts no change with respect to control (untreated) parasites. X axis depicts log2 fold change and Y axis shows −log10 p values. The scatter plot represents the expression level of each gene under control (untreated) and E2G12 treated condition. Each dot represents a gene. The vertical position of each gene represents its expression level in the E2G12 treated sample while the horizontal position represents its expression level in the control (untreated) samples. These genes that fall above the diagonal are over expressed (red dots) and genes that fall below the diagonal are downregulated (green dot) as compared to their median expression level in experimental grouping. Black dots represent no change in the expression level. X axis represents control (untreated) and Y axis represents E2G12 treated **(E)** Heatmap of 50 most differentially expressed *P. falciparum* 3D7 mRNAs at trophozoite stage after the rescue from E2G12 treatment. Gene IDs were identified in plasmodb.org data base. The scale of the mRNA expression level was from 0 to 6, where 0 was comparable to control (untreated) parasites and 6 depicts the highest expression of the transcript after rescue. **(F)** Quantitative analysis of *P. falciparum* 3D7 mRNA transcripts by log2 fold change after the rescue from E2G12 treatment at trophozoite stage. The upregulated and downregulated transcripts with gene IDs were depicted on a scale of log2 fold change from −2 to +8 on Y axis. mRNA transcripts were analysed in biological replicates N=3. **(G)** Volcano plot showing the changes in total mRNA transcripts after the rescue from E2G12 treatment. Each dot signifies a mRNA transcript. Green depicts downregulation of mRNA level; red depicts upregulation of mRNA and black depicts no change with respect control (untreated) parasites. X axis depicts log2 fold change and Y axis shows −log10 p values. **(H)** The scatter plot represents the expression level of each gene under E2G12 treated and rescue conditions. Each dot represents a gene, red, green and black signifies upregulated, downregulated and no change transcripts levels. **(I)** Heatmap of 50 most differentially expressed *P. falciparum* NF54 mRNAs at trophozoite stage after E2G12 treatment. Gene IDs were identified in plasmodb.org data base. The scale of the mRNA expression level was from 0 to 2, where 0 was comparable to control (untreated) parasites and 2 depicts the highest expression of the transcript. **(J)** Quantitative analysis of *P. falciparum* NF54 mRNA transcripts by log2 fold change after E2G12 treatment at trophozoite stage. The upregulated and downregulated transcripts after E2G12 treatment were depicted in X axis on a scale of log2 fold change mRNA transcripts were analysed on biological replicates N=3. **(K)** Volcano plot showing the changes in total mRNA transcripts after the treatment with E2G12 in *P. falciparum* NF54 parasites. Each dot signifies a mRNA transcript. Green depicts downregulation of mRNA level; red depicts upregulation of mRNA and black depicts no change with respect control (untreated) parasites. X axis depicts log2 fold change and Y axis shows −log10 p values. **(L)** The scatter plot represents the expression level of each gene under E2G12 treated and control (untreated) conditions. Each dot represents a gene, red, green and black signifies upregulated, downregulated and no change transcripts levels respectively. **(M)** Relative abundance of mRNA transcripts of *P. falciparum* 3D7 and NF54 parasites at trophozoite stage after E2G12 treatment and antibody rescued conditions. Total 6 transcripts were analysed having biological replicates N=3. Statistical P values were denoted as *. P<0.05 as *, P<0.01 as **, P<0.001 as *** and P<0.0001 as ****.

Interestingly, upon rescue from FAs import inhibition and concomitant with the progression of parasite nuclear division and H2A, H2B, H3, and H4 rescue, ClpM and SufB transcripts were downregulated by log2 1-2-fold (Figure. 3E & 3F, Excel sheet. S6). Upon rescue, several genes were upregulated by log2 3-4-fold to rescue the parasite physiology, which include ribosomal protein P2 (PfP2), acyl CoA-synthetase, Patatin-like phospholipase 1-4, dynein Heavy Chain (DHC), Protein Kinase 6 (PK6), Enoyl acyl carrier reductase (Figure. 3F, Excel sheet. S6). After E2G12 treatment, histone transcripts were downregulated whereas upon rescue H2A, H2B, H3 and H4 all were upregulated (Figure. 3F, Excel sheet. S5 & S6). Heatmap and log2 fold change data of these genes after E2G12 treatment and subsequent rescue seemed to suggest a direct correlation of cell cycle regulation with the FAs import and through TAG formation and downstream membrane biogenesis through the Kennedy pathway. Upon rescue from E2G12 treatment, overall, a significant upregulation of transcripts was observed as represented by the volcano and scatter plot (Figure. 3G & 3H, Excel sheet. S6). Here it is important to note that in FAs import inhibited parasites, the transcript level of PfP2 did not change with respect to untreated parasites but significantly upregulated in rescued parasites (Figure. 3M, Excel sheet. S5 & S6), which suggested that after rescue, PfP2 protein expression was urgently required to establish the FAs importing complex on the iRBC surface and import FAs to mitigate the scarcity in the parasites and proceed for membrane biogenesis. Since we did not observe PfP2 transcript level difference in E2G12 treated parasites, we chose not to use PfP2-3xHA-glmS transgenic line to degrade PfP2 mRNA using glucosamine (GlcN) and send parasites for transcriptomics analysis, rather we used E2G12 and blocked FAs import to analyse fold change of transcripts and narrow down the pathway through which histone degradation was achieved to arrest nuclear and cell division. In *P. falciparum* 3D7, the relative abundance of ClpP transcript was downregulated after E2G12 treatment, but significantly upregulated in rescued parasites whereas ClpR was upregulated in E2G12 treated parasites and upon rescue the transcript level went up further (Figure. 4C). But in *P. falciparum* NF54, the transcript level of ClpP and ClpR both went up at PAI 34h concomitant with the restoration of histone H3 and H4 expression (Figure. 1F & 1H and Figure. 4C, Excel sheet. S2). The function of ClpM is to hydrolyse ATP and generate energy for the functioning ClpP/ClpR machinery ^45, 46^. The transcript level of ClpM was significantly upregulated in E2G12 treated 3D7 and NF54 parasites, and upon rescue from the E2G12 treatment, ClpM transcript went down. This observation suggested that when ClpP/ClpR machinery was in action, the ATP hydrolysis by ClpM was necessary to carry out the required function. Unlike ClpP/ClpR/ClpM, the transcript level of E3 ubiquitin ligase, Proteasome subunit RPN1 and RPN6 did not change in the E2G12 treated *P. falciparum* 3D7 parasites, which suggested that histone degradation possibly was not through the ubiquitination and proteasome-mediated (Figure. 3M, Excel sheet. S5). On the contrary, in the rescued *P. falciparum* 3D7, the transcript level of E3 ubiquitin ligase, Proteasome subunit RPN1, and RPN6 were significantly upregulated (Figure. 3M, Excel sheet. S6). This seemed to suggest that, due to the arrest, if there were misfolded proteins, they needed to be degraded through ubiquitin mediated proteasome machinery to clean the system from junk proteins and make the system get going and proceed towards cell division.

**Figure 4.**
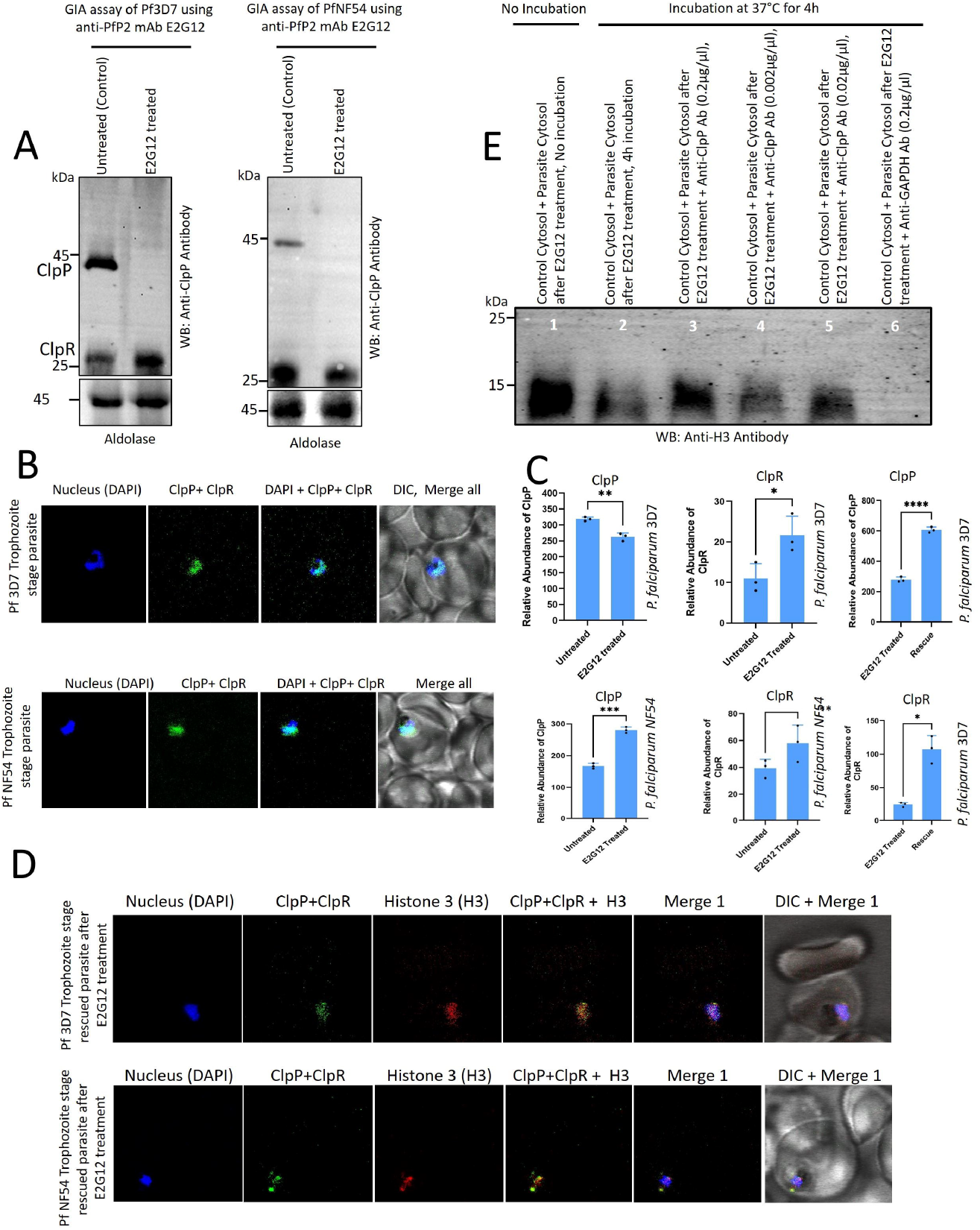
Evolutionary conserved bacterial ClpP/ClpR protein quality control machinery degrades histones in the *P. falciparum* parasite nucleus. **(A)** Immunoblot of *P. falciparum* 3D7 and NF54 trophozoite stage parasites under Control (untreated) and E2G12 (700 ng/µl) treated GIA condition at 37°C, PAI 24h. Blot was probed with Anti-ClpP rabbit mAb. Parasite aldolase was used as a loading control. **(B)** Immunofluorescence Assay (IFA) of control *P. falciparum* 3D7 and NF54 trophozoite stage parasites stained with Anti-ClpP rabbit mAb. Anti-rabbit Alexa488 conjugated secondary antibody was used for fluorescence. Parasite nucleus was stained with DAPI. **(C)** Relative abundance of ClpP and ClpR transcripts in *P. falciparum* 3D7 and NF54 trophozoite stage parasites under control (untreated) and E2G12 treated condition. Statistical P values were denoted as *. P<0.05 as *, P<0.01 as **, P<0.001 as *** and P<0.0001 as ****. **(D)** Immunofluorescence Assay (IFA) of E2G12 treated *P. falciparum* 3D7 and NF54 trophozoite stage parasites. Parasites were stained with Anti-ClpP rabbit mAb and Anti-Histone 3 (H3) mouse mAb. Appropriate anti-rabbit and anti-mouse Alexa 488 and Alexa 647 were used. Parasite nucleus was stained with DAPI. **(E)** Cross-incubation of *P. falciparum* 3D7 trophozoite stage cytosol of control (untreated) and the cytosol of E2G12 treated parasites without protease inhibitor. In lane 3-5, E2G12 treated parasite cytosol was initially treated with anti-ClpP mAb for 2h at 37°C and lane 6 was treated with GAPDH antibody. Anti-ClpP mAb treated E2G12 parasite cytosol was then incubated with control parasite cytosol for 4h at 37°C. Lane 1, as a positive control, control parasite cytosol was mixed with E2G12 treated parasite cytosol and immediately denatured using loading dye. From lane 2-6, all mixtures were incubated for 4h at 37°C. Immunoblot was probed with anti-Histone 3 (H3) antibody. Anti-rabbit HRP tag secondary antibody was used for ECL imaging.

When FAs import was inhibited using E2G12 antibody, the level of Triacylglycerol (TAG) was significantly enhanced in the parasites ^12^. Cellular TAG homeostasis is the only metabolic process known which is directly regulated by Cdk1/Cdc28-dependent phosphorylation of key anabolic and catabolic enzymes such as Tgl4, which suggests the importance of TAG and its continuous mobilization during the cell cycle progression ^47^. The FAs and membrane perturbation induce G1 cell cycle arrest, confirming the importance of membrane biogenesis for cell cycle progression ^48^. Comparison between E2G12 treated and E2G12 rescued parasites revealed that Tgl4 homologue in *P. falciparum* 3D7 might be Exported Lipase 1 / 2 (XL1/XL2), PF3D7_1001400 / PF3D7_1001600 or Patatin-like phospholipase 1 (PATPL1) PF3D7_0209100. XL1 / XL2 or PATPL1 might be involved in the regulation of TAG levels in the parasites. In E2G12 rescued parasites in comparison with arrested parasites, the XL1 and PATPL1 gene expression was upregulated by log2 4-5-fold (Figure. S3E & S3F and Excel sheet. S5 & S6). To cross verify the involvement of XL1 / XL2 in the TAG regulation, parasites were treated with Orlistat, which blocks the conversion of TAG to DAG, hence TAG level increases ^10^. We wanted to check the expression level of XL1 / XL2 in Orlistat treated parasites. To our surprise, we did see a significant change in the expression level of XL1 and PATPL1 in Orlistat treated parasites whereas, XL2 and PATPL4 did not show any difference (Figure. S3, Excel sheet. S7). So, the pathway following which the TAG levels was increased determines which enzyme should be in charge of regulating the TAG level in the parasites. When TAG level was increased due to FAs import inhibition through the blockage of PfP2 on the iRBC surface ^12^, XL1 / PATPL1 appeared to regulate TAG level in the parasites. In the E2G12 treated FAs import inhibited parasites, where TAG level was high, the H2A, H2B, H3, and H4 was degraded. But, when TAG level was increased by Orlistat, we did not see H2A, H2B, H3 and H4 degradation (Figure. 1D). It suggested a dichotomy, where it indicated that merely enhanced TAG may not have a direct correlation with the degradation of H2A, H2B, H3 and H4 rather it was a pathway specific response and depends on the mechanism as to how TAG level was enhanced and how it is being regulated with the progression of membrane biogenesis and cell cycle progression.

Tgl4 is a triacylglycerol (TAG) lipase in *Saccharomyces cerevisiae* ^47^. Tgl4 catalyzes the breakdown of TAG, which are stored as energy reserves in lipid droplets ^10,47^. Tgl4 is activated by phosphorylation by the cell-cycle kinase Cdk1/Cdc28 during the G1/S transition, which allows yeast to enter the cell cycle from a stationary phase ^47^. To understand the molecular regulation of TAG level and its correlation with cell cycle regulation through Cyclin Dependent Kinase (CDK6) / Protein Kinase 6 (PK6) and Cyclin Dependent Kinase (CDK9) / Protein Kinase 9 (PK9), transgenic PK6-2xHA and PK9-2xHA parasites were used for immunoprecipitation (IP) assay to check whether XL1/PATPL1 interacts with PK6 and/or PK9 in *P. falciparum* 3D7 transgenic parasites (Figure. S3, Excel sheet. S8). Transgenic PK6 and PK9 appeared to be a 45 kDa protein in western blot using anti-HA antibody (Figure. S3A). IP of PK6 and PK9 were done, and the interactor proteins were analyzed as per the mass spectrometry strategy (Figure. S3B). PK6 and PK9, both were found to be interacting with PATPL1 but not with XL1 (Figure. S3C & S3D). In addition, cell division-related other proteins were also found to be interacting, which suggested the authenticity of the observation and established the role of PK6/PK9 in the regulation of PATPL1. The PK6 transcript level in E2G12 treated cell cycle arrested *P. falciparum* 3D7 parasites was comparable to that of untreated parasites and significantly upregulated in the E2G12 rescued parasites (Figure. 3M, Excel sheet. S5 & S6). This suggested that the activity of PATPL1 may be regulated through phosphorylation by PK6. When PK6 level is low, the required phosphorylation of PATPL1 was not achieved, hence PATPL1 became inactive and did not catabolize TAG to DAG and FAs for PC biosynthesis through the Kennedy pathway ^38^. Now, in the rescued parasites, PK6 transcript level was higher hence PATPL1 phosphorylation was achieved and subsequently TAG was catabolized to FAs for PC biosynthesis and membrane biogenesis. Phylogenetic tree analysis of PATPL1-4 and several other Lipase enzymes in humans and in yeast seemed to suggest that PATPL1 existed in a different branch, deviating from other lipase and might have some unique functions in the malaria parasites related to TAG level regulation and maintaining membrane homeostasis and bi-layer assembly (Figure. S3G).

FAs import inhibition in *P. falciparum* NF54 parasites led to the male and female gametocyte formation (Figure. 2A). After E2G12 treatment at PAI 34h (Post Antibody Incubation) total transcriptomics revealed the upregulation of *de novo* FAs biosynthesis-related genes in the apicoplast, in addition, iron Sulphur cluster assembly protein SufB and histone transcripts were also upregulated (Figure. 3I, 3J, Excel sheet. S2). This corroborated with the observation in Figure. 1F & 1H. The *de novo* FAs biosynthesis related transcripts FabB/FabF (PF3D7_0626300), FabZ (PF3D7_1323000), KASIII (PF3D7_0211400) in *P. falciparum* NF54 were significantly upregulated (Figure. 3I & 3J, Excel sheet. S2). Interestingly, the master regulator for gametocyte induction AP2-G transcript were upregulated by log2 2-3-fold, which suggested the reason for gametocyte induction and a possible mechanism through FAs import (Figure. 3I, Excel sheet. S2). In FAs import inhibited *P. falciparum* NF54 parasites, transcription activator Tat binding protein 1 (TBP-1) and transcriptional co-repressor HCNGP were upregulated by log2 3-4-fold (Figure. 3J, Excel sheet. S2). In the apicoplast, DNA-directed RNA polymerase subunit β (PF3D7_API04200/ PF3D7_API04300/ PF3D7_API04400) was upregulated by log2 3-4-fold (Figure. 3J, Excel sheet. S2). Inhibition of FAs import led to a significant upregulation of genes in NF54 parasites as compared to downregulation and interestingly, a large number of genes were also unaltered (Figure. 3K & 3L, Excel sheet. S2). All these transcriptional changes suggested a drastic molecular level change which led to the initiation of *de novo* biosynthesis of FAs, induction and the formation of male and female gametocytes and a changed lipid homeostasis favouring gametocyte membrane biogenesis and bi-layer assembly.

FAs import inhibition led to a significant upregulation and downregulation of many genes, resulted into histone degradation and cell cycle arrest of malaria parasites. We wondered how the cross-talk between FAs import, membrane biogenesis, gametocyte induction and cell cycle regulation through transcript regulation was achieved and whether this transcription regulation of individual genes was somehow directed by Long non-coding RNAs (LncRNAs) at trophozoite stage of the parasites. To assess this hypothesis, we identified and quantified the entire population of LncRNAs (now it will be called as PfLncRNAoms) in *P. falciparum* 3D7 and NF54 at trophozoite stage after E2G12 treatment and under E2G12 rescued conditions (Figure. S4A-S4H and Figure. S5A-S5D, Excel sheet, S10, S11 & S12). Under FAs import inhibited and cell cycle rescued condition, in the PfLncRNAoms of *P. falciparum* 3D7 and NF54 parasites, many LncRNAs responded to the FAs deprived condition and showed a significant upregulation and downregulation of LncRNAs from the identified chromosome under treated and E2G12 rescued condition (Figure. S4I, S4J & S5E). To assess the direct corelation of an individual LncRNA with the E2G12 treated (FAs deprived condition) and E2G12 rescued condition (cell cycle rescued condition), when transcript level of LncRNA was quantified, it was observed to be responding to the rescued condition and consequentially the expression level of upregulated LncRNAs came back under rescued condition (Figure. S4B & S4F). In essence, each LncRNA responded to the FAs deprived condition and membrane biogenesis by changing its expression under treated and rescued condition, which suggested a direct role of PfLncRNAoms in the regulation of a set of genes responsible to control parasite physiology having to do with parasite membrane biogenesis, FAs biosynthesis, DNA replication and cell cycle regulation.

### Bacterial ClpP/ClpR serine protease in the parasite nucleus degrades histones

In *P. falciparum* 3D7 and NF54 parasites, ClpP (PF3D7_0307400) and ClpR (PF3D7_1436800) both appeared to be essential ^49, 50.^ Both the proteins localized in the parasite nucleus having ClpP molecular wt. 43 kDa and ClpR 28 kDa (Figure. 4A). In the FAs import inhibited parasites, astonishingly, the expression level of 43 kDa band of ClpP was very low as compared to 28 kDa ClpR (Figure. 4A). Rather, the expression level of 28 kDa band seemed to be upregulated in *P. falciparum* 3D7 parasites but not in *P. falciparum* NF54 (Figure. 4A). This raised a legitimate question as to why ClpP expression downregulated in the E2G12 treated parasites? On the flip side, this observation suggested that the degradation of H2A, H2B, H3, and H4 may be carried out by the ClpR and not by ClpP or may be in combination of both. The ClpP and ClpR appeared to localize in the parasite nucleus in *P. falciparum* 3D7 and NF54 (Figure. 4B). However, since the Anti-ClpP antibody also interacts with ClpR, it is still a question whether nuclear localization of ClpP was due to both or either of that? Corroborating with the Western blot data in Figure. 4A, the transcript level of ClpP in *P. falciparum* 3D7 was significantly downregulated after E2G12 treatment but in contrary, in *P. falciparum* NF54 at PAI 34h, the ClpP transcript was significantly upregulated (Figure. 4C, Excel sheet. S2 & S5). The transcript level of ClpR in the *P. falciparum* 3D7 and NF54 was significantly upregulated (Figure. 4C, Excel sheet. S2 & S5). It was clearly evident that the FAs import inhibition directly affected the transcription and translation regulation of ClpP and ClpR in the parasite nucleus, which might have downstream functional implications in the regulation of parasite cell division. In the rescued *P. falciparum* 3D7 parasites after E2G12 treatment, ClpP and ClpR colocalized with rescued histone 3 (H3) in the parasite nucleus (Figure. 4D), which further confirmed that upon E2G12 removal from the culture medium under GIA condition, the expression level of H3 came back and ClpP and ClpR both localized in the parasite nucleus suggesting a critical role of ClpP and ClpR in the quality control and turnover of histones during cell cycle progression and regulation. In untreated HEK239T mammalian cells, ClpP showed localization mostly in the cell cytoplasm and did not colocalize with the nucleus (Figure. S1C). However, after Cerulenin and C75 treatment, the nucleus of some HEK293T cells appeared to stain with ClpP antibody, which seemed to suggest a possible translocation of cytoplasmic ClpP into the nucleus. But the majority of cells showed cytoplasmic ClpP localization after Cerulenin and C75 treatment (Figure. S1C). In *E. coli*, ClpP stained as intense green puncta in the bacterial cell cytosol (Figure. S6C). Interestingly, there was heterogeneity in the number of green puncta; some bacteria showed one and some showed more than one in a single bacterial cell (Figure. S6C). In *Cyanobacteria*, ClpP was stained throughout the bacterial cytoplasm, but interestingly, it appeared that dividing *Cyanobacteria* had more intense staining as compared to other cells, suggesting an upregulated expression of ClpP at certain stages of the cell cycle (Figure. S6E).

Genetic knockout study seemed to suggest that ClpP and ClpR are essential genes, which indicated their critical role in parasite physiology^49,50^. Our study suggested an important role of ClpP and ClpR in the regulation of parasite cell cycle through histone’s quality control and turnover. Our study showed why ClpP and ClpR genes were indispensable for parasite development. In addition. ClpP and ClpR might have additional functions upstream of histone quality control and hence knocking out or conditional downregulation of both genes might affect parasite physiology and arrest the development which may not have direct corelation with the parasite cell cycle. Under this context, we chose not to go for genetic knockdown of ClpP and or ClpR using a conditional downregulation system e.g., ribozyme-glmS or tetR-DOZI aptamer, to show a direct role of ClpP and ClpR on the degradation of histones. Instead, we took a completely different approach using anti-ClpP antibody to confirm the direct role of ClpP and ClpR in the degradation of histones when FAs import was inhibited. When FAs deprived parasite cytosol was cross incubated with the control parasite cytosol, the histones in the control parasites were degraded (Figure. 1K & 1L). To confirm whether ClpP and ClpR are directly involved in this degradation of histones, we cross incubated control parasite cytosol with FAs import inhibited parasite cytosol in the presence of varying concentration of anti-ClpP antibody. FAs import inhibited parasite cytosol was first pre-incubated with anti-ClpP antibody to block the function of ClpP and ClpR and then cross-incubated with untreated parasite cytosol containing histones. As a positive control to check the degradation, again control parasite cytosol was cross incubated with FAs import inhibited parasite cytosol (Figure. 1K). We used GAPDH antibody to rule out the effect of stearic hindrance due to the presence of antibody molecules on the ability of degradation by ClpP and ClpR in the solution. This entire permutation and combination strategy strongly suggested that when the function ClpP and ClpR was blocked by the presence of anti-ClpP antibody, histone was largely not degraded depending on the concentration of the anti-ClpP antibody (Figure. 4E). Whereas, the GAPDH antibody could not resist the degradation of histones by ClpP and ClpR. This experimental jugglery confirmed that ClpP and ClpR are directly involved in the degradation of histones, which may be for quality control and turn over during cell cycle progression.

### Hyperphosphorylation of arginine in histones marks for degradation by ClpP/ClpR

In a biological cell, histone is the only protein that gets heavily loaded with post translationally modified (PTMs) amino acids ^51, 52, 53, 54, 55, 56, 57^. The PTMs in histones are known to be absolutely essential in transcription regulation, DNA replication, nucleosome formation, genome topology, DNA packaging through condensation and de-condensation and in various cell cycle and tissue differentiation-related biological activities and beyond^51, 52, 53, 54, 55, 56, 57^. We hypothesised that FAs import inhibition led to unique PTMs in the parasite histones, which compromised the quality and, as a result, marked for removal through degradation by the nuclear localized ClpP/ClpR protein quality control machinery. To pursue this daunting hypothesis, we did not go for selective isolation of histone species using antibodies, rather, we isolated total histones using acidified ethanol (Ac.EtOH) ^58^ as per the strategy discussed (Figure. 5A). Ac.EtOH extracted histones were conformationally active as anti-H3 and anti-H4 antibody binding was confirmed in western blotting (Figure. 5B & 5C). This established the authenticity of the isolation process and maintenance of the quality of histones. Additionally, we were curious to check whether Ac.EtOH extracted histones still retain the ability to be degraded by ClpP/ClpR machinery. When Ac.EtOH extracted histones were cross-incubated with E2G12 treated (FAs import inhibited) parasite cytosol, the H3 and H4 band intensity significantly reduced in western blotting (Figure. 5D & 5E), which suggested that ClpP/ClpR machinery was still active on the Ac.EtOH extracted histones possibly because of conformational integrity and maintenance of histone’s PTMs after Ac.EtOH extraction.

**Figure 5.**
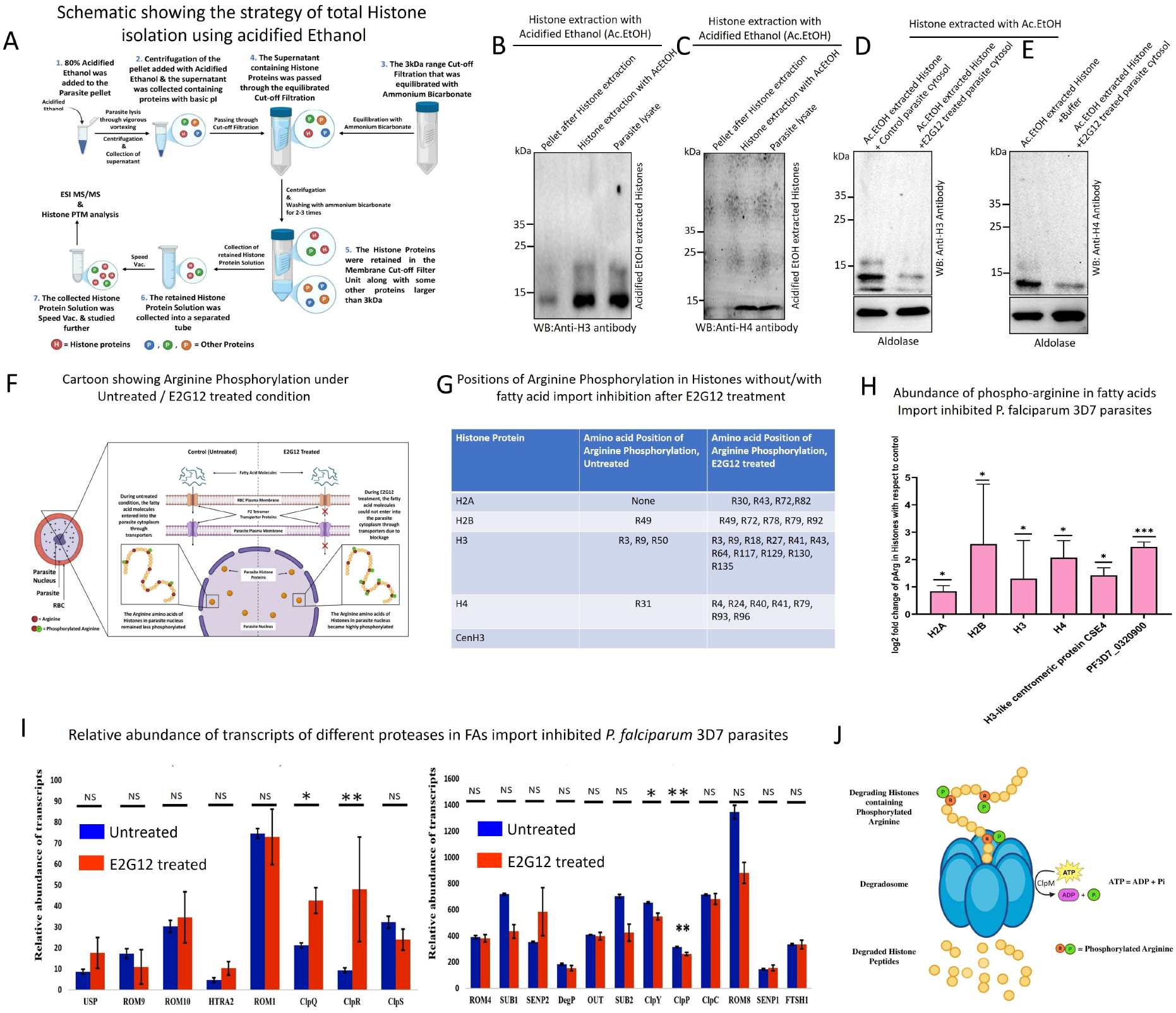
Arginine hyperphosphorylation by arginine kinase is a prerequisite step for the action of ClpP/ClpR machinery in the parasite nucleus. **(A)** Schematic showing the strategy of total histone isolation using acidified ethanol (Ac.EtOH). **(B-C)** Immunoblot of histones before and after extraction with Ac.EtOH. The blot was probed with anti-H3 and anti-H4 antibody. **(D-E)** Cross-incubation of Ac.EtOH extracted histones with control and E2G12 treated parasite cytosol without protease inhibitor. Incubation was carried out at 37°C for 4h. Immunoblot was probed with anti-H3 and anti-H4 antibody. Parasite aldolase was used as a loading control. **(F)** Carton showing histone hyperphosphorylation in the parasite nucleus under control (untreated) and E2G12 treated condition **(G)** Table showing the positions of arginine phosphorylation in histones H2A, H2B, H3 and H4 in *P. falciparum* 3D7 trophozoite stage control (untreated) and E2G12 treated parasites. Positions of arginine phosphorylation in histones were identified by qTOF MS/MS mass spectrometry facility. **(H)** log2 fold change of the abundance of phospho-arginine (P-Arg) in histones, in H3 like centromeric protein CSE4 and hypothetical protein PF3D7_0320900. **(I)** Relative abundance of transcripts of 20 different proteases in P. falciparum 3D7 trophozoites stage parasites under untreated (control) and E2G12 treated conditions. **(J)** Schematic showing the possible mechanism of histone degradation by ClpP/ClpR machinery after hyperphosphorylation of arginine residues. Statistical P values were denoted as *. P<0.05 as *, P<0.01 as **, P<0.001 as *** and P<0.0001 as ****.

In bacteria, arginine phosphorylation (P-Arg) by arginine kinase McsB marks proteins for degradation under a protein quality control process by ClpP machinery ^59, 60, 61, 62, 63^. We hypothesized that FAs import inhibition and compromised membrane biogenesis led to hyperphosphorylation of arginine in histones as described in the carton (Figure. 5F). Hyperphosphorylation of arginine marked histones for degradation by ClpP/ClpR machinery in the parasite nucleus. To check the extent of P-Arg residues in histones after Ac.EtOH extraction from FAs import inhibited parasites, we performed qTOF mass spectrometry based PTMs analysis of histones and identified which arginine was phosphorylated and their position in H2A, H2B, H3 and H4 (Figure. 5G). In the untreated control parasites, H2A did not show any P-Arg residues, whereas, H2B was identified to have one P-Arg residue at R49, H3 had three P-Arg residues at R3, R9 and R50 and H4 had one P-Arg residue at R31. In the FAs import inhibited parasites, H2A, H2B, H3 and H4 all were heavily modified and identified with multiple P-Arg residues at multiple positions (Figure. 5G, Excel sheet. S13). The abundance of P-Arg was significantly upregulated by log2 2-3-fold in all types of histones (Figure. 5H). This suggested about a cellular mechanism that became active under FAs deprived condition having the unique ability to phosphorylate arginine residues in histones and marked for degradation through ClpP/ClpR machinery in the parasite nucleus.

Thereafter, we wondered is there any arginine kinase or Creatine kinase enzyme in Plasmodium, which has the ability to phosphorylate arginine and might have homology with the bacterial McsB and Toxoplasma. Surprisingly, there is no homology of McsB in Plasmodium. But in Toxoplasma, ATP: guanido phosphotransferase (TGME49_500283), a putative arginine kinase was 27-30% similar to McsB. On the contrary, in Plasmodium, there is no homolog of ATP: guanido phosphotransferase. Hence, we wondered about the identity of an unknown arginine kinase or creatine kinase in Plasmodium and its function. In invertebrates and some microbes, arginine kinase acts as an energy buffer by reversibly transferring a phosphate group from ATP to L-arginine, hence, creating high energy P-Arg to regenerate ATP during instantaneous need^60, 61, 62, 63^. Whether P-Arg donates high energy PO_4_^3-^ for ATP generation during rapid asynchronous schizogony as an alternative source of energy is not yet looked at. Experimentally, to answer a specific question, we took a promiscuous approach by using a polyclonal antibody to arginine kinase of Honeybee (*Apis mellifera*) and polyclonal antibody to creatine Kinase of human and tried to find out whether any protein / enzyme in Plasmodium interacts with the arginine kinase antibody or creatine Kinase antibody presumably retaining the antibody binding domain and preserving the arginine kinase or creatine kinase activity. Arginine kinase antibody interacted with a protein in Plasmodium showed~22-23 kDa molecular weight (Figure. 6A). This protein band appeared in the untreated (control) and E2G12 treated parasites but was very low in its expression in the E2G12 rescued parasites (Figure. 6A). Interestingly, the expression level of this unknown arginine kinase did not change when the cell cycle of the parasite was arrested by Taxol, Aphidicolin and Orlistat (Figure. 6B). The arginine phosphorylation by this unknown arginine kinase seemed to be responding to the E2G12 rescued condition where the arginine kinase expression level was down may be to accelerate histone production, nucleosome formation and cell cycle progression. The arginine phosphorylation by the novel arginine kinase corroborated with the P-Arg PTM analysis in the histones of untreated (control), and E2G12 treated parasites (FAs import inhibited condition) (Figure. 5F, 5G & 5H). Interestingly, in the untreated parasite lane, we also observed the expression of arginine kinase but the histones of untreated parasite showed few phosphorylated arginine residues (Figure. 6A & 5G). This seemed to suggest a protection mechanism that shields arginine residues from being phosphorylated or a role of a phosphatase that maintains the balance of arginine phosphorylation and dephosphorylation under normal parasite physiology. Immunoblot analysis of bacterial ClpP, arginine kinase and creatine kinase in *E. coli* and *Cyanobacteria* showed distinct bands confirming their presence in the prokaryotic system and antibody specificity (Figure. S5A). IFA of *E. coli* and Cyanobacteria showed the localization of arginine kinase and creatine kinase in the cell cytoplasm (Figure. S6D, S6F, S6G & S6H). The distribution of arginine kinase in Cyanobacteria appeared to be homogeneous throughout the cytoplasm however, few cells showed a higher intensity of staining which may be due to the differences in cell cycle stages (Figure. S6F).

**Figure 6.**
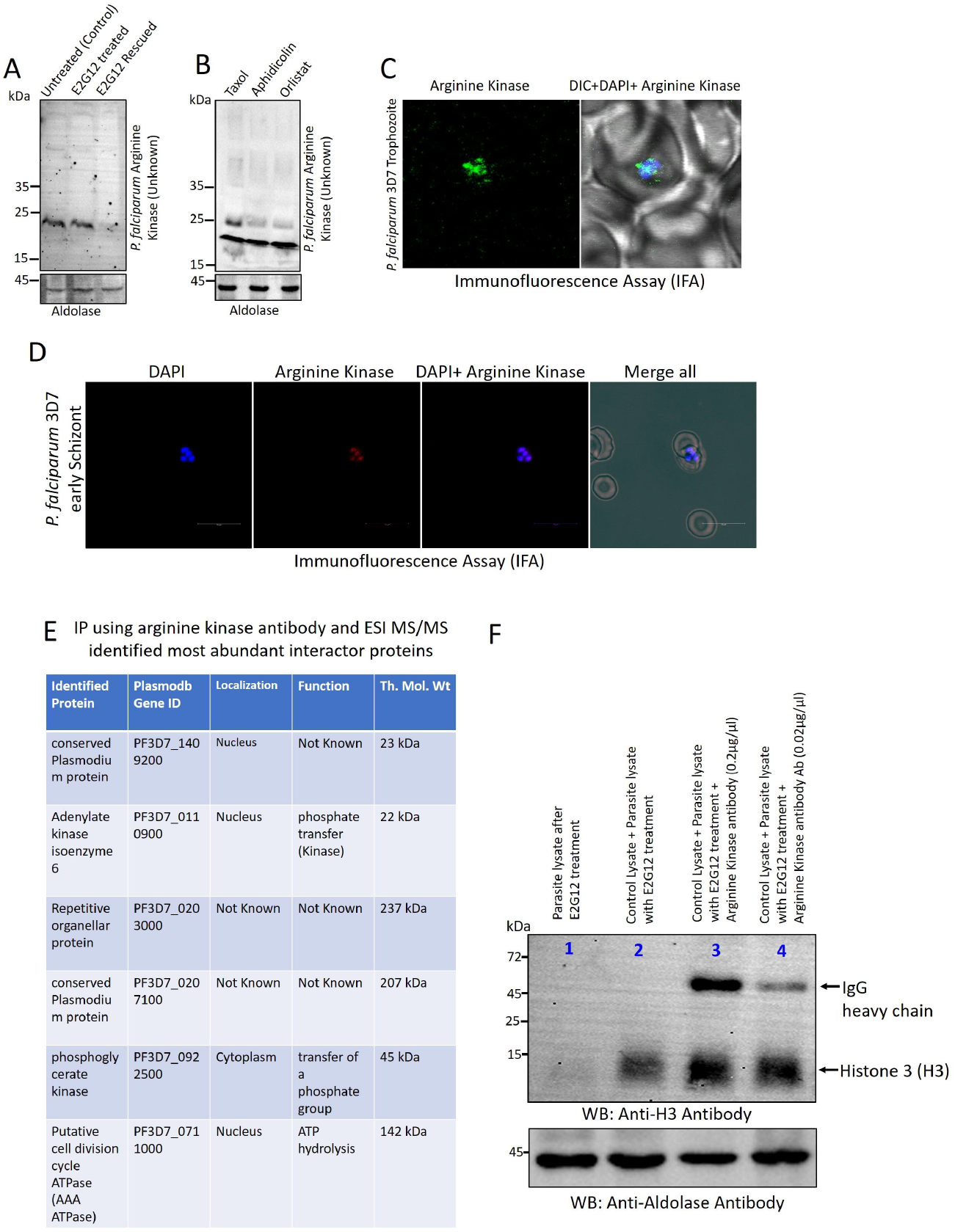
Hypothetical protein (PF3D7_1409200) possibly acting as an arginine kinase adding PO_*4*_^*3−*^ to the arginine residues in histones. **(A)** Immunoblot of untreated (control), E2G12 treated and E2G12 rescued *P. falciparum* 3D7 parasites probed with arginine kinase antibody which was developed against Honeybee (*Apis mellifera*) arginine kinase. Parasite aldolase was used as a loading control. **(B)** Immunoblot of Taxol, Aphidicolin and Orlistat treated *P. falciparum* 3D7 trophozoite stage parasites probed with arginine kinase antibody. Aldolase was used as a loading control. **(C & D)** IFA of *P. falciparum* 3D7 trophozoite and early schizont stage parasites showing nuclear localization of arginine kinase **(E)** Table showing interactors proteins after Immunoprecipitation (IP) using arginine kinase antibody and qTOF ESI MS/MS mass spectrometry-based identification of arginine kinase and its interactor proteins. **(F)** Cross-incubation of *P. falciparum* 3D7 trophozoite stage cytosol of control (untreated) and the cytosol of E2G12 treated parasites without protease inhibitor. In lane 1, E2G12 treated parasite cytosol was loaded, in lane 2, control parasite cytosol was cross-incubated with E2G12 parasite cytosol and in lane 3-4, control parasite cytosol was initially treated with varying concentration (0.2µg/µl and 0.02µg/µl) of anti-arginine kinase antibody for 2h at 37°C and then incubated with E2G12 treated parasite cytosol. After arginine kinase antibody treatment, E2G12 parasite cytosol was then incubated for 4h at 37°C. Immunoblot was probed with anti-Histone 3 (H3) antibody. Anti-Aldolase antibody was used as a loading control.

IFA using arginine kinase antibody localized the enzyme in the *P. falciparum* parasite nucleus and showed colocalization with nuclear stain DAPI (Figure. 6C & 6D). Thereafter, we wondered about the identity of this unknown arginine kinase. Immunoprecipitation assay (IP) using arginine kinase antibody and qTOF mass spectrometry of E2G12 treated *P. falciparum* 3D7 trophozoite stage parasites identified the gene of arginine kinase as a hypothetical protein having plasmodb gene ID PF3D7_1409200 (Figure. 6E, Excel sheet. S14). This was the most abundant protein in the IP sample. In addition, adenylate kinase like gene (PF3D7_0110900), was also identified (Figure. 6E, Excel sheet. S14). Possibly, the arginine kinase activity turned out to be carried out by this hypothetical protein (PF3D7_1409200) having a nuclear localization signal and adenylate kinase isoform 6 (PF3D7_0110900) in the parasite nucleus may be involved as a PO_4_^3-^ donor. The molecular weight of both the identified proteins was ~ 22-23 kDa as identified in western blot data (Figure. 6A & 6B). The function of this hypothetical gene (PF3D7_1409200) in Plasmodium was never identified before hence this is the first report describing the probable function of a new arginine kinase like gene. Henceforth, we would like to call this gene, PF3D7_1409200 as *NPD1951-AK* in the plasmodium genome database. After the identification of the *NPD1951-AK gene*, we wanted to check the direct involvement of the kinase activity of *NPD1951-AK* in the arginine hyperphosphorylation of histones and as a result the activation of histone degradation by ClpP/ClpR machinery. To understand the importance of arginine phosphorylation in order for ClpP/ClpR activity to achieve histone degradation, we blocked the phosphorylation of histones by inhibiting the activity of *NPD1951-AK* by dose dependent incubation with arginine kinase antibody in the E2G12 treated parasite cytosol. Thereafter, we questioned whether blocked *NPD1951-AK* has any consequences on the activity of ClpP/ClpR when incubated with the untreated (control) parasite cytosol. Interestingly, when arginine kinase antibody was used to block the activity of *NPD1951-AK*, we did not see histone H3 degradation (Figure. 6F). At two arginine kinase antibody concentrations, 0.2µg/µl and 0.02 µl/µl, we observed reduced H3 degradation. This could possibly mean that due to the lack of phosphorylation of histones, the H3 degradation was inefficient by ClpP/ClpR machinery and also depicted the importance of phosphorylation for ClpP/ClpR activity. In the identified interactor of *NPD1951-AK*, adenylate kinase like protein (ADKLP) (PF3D7_0110900) was another abundant protein having a predicted nuclear localization signal (Figure. 6E & S8H). The homology of ADKLP was significantly similar to human adenylate kinase isoform 6, which is known to localize in the cell nucleus ^71. 72, 73^. Adenylate kinase is a phosphotransferase that maintains cellular energy homeostasis by catalysing the reversible reaction 2ADP⇌ATP+AMP ^71, 72, 73^. The importance of ADKLP in the activity of *NPD1951-AK* is not clear and the sources of PO_4_^3-^ for arginine phosphorylation is not yet known.

Thereafter, we questioned whether, other than Clp family of serine proteases, was there any other types of proteases which got upregulated under FAs import inhibited condition and the possibility of protein degradation by those proteases. To address this question, we measured the relative abundance of transcripts of 20 different proteases under untreated Vs E2G12 treated condition in *P. falciparum* 3D7 parasites. Except ClpP, ClpR, ClpQ and ClpY, the transcript level of 16 other proteases did not change significantly (Figure. 5I, Excel sheet. S5), which indicated that FAs import inhibition did not cause another protease to upregulate. Rather it was selectively changed the expression pattern of Clp family of proteases, which implied a direct mechanism of activation of Clp proteases under the FAs deprived condition. Our data suggested that FAs import inhibition and compromised membrane biogenesis led to hyperphosphorylation of arginine residues in histones. The uncanny level of P-Arg led to compromised quality of histones which marked for degradation through ClpP/ClpR machinery (Figure. 5J). Histone degradation plausibly led to the downstream arrest of DNA replication and cessation of histone octamer containing nucleosome formation hence, causing cell cycle arrest until the upstream FAs import problem was fixed and membrane biogenesis was resumed for nuclear membrane assembly.

## Discussion

Our study has discovered a novel cell cycle checkpoint, Lipid (L) checkpoint, showing a crosstalk mechanism between FAs import or *de novo* biosynthesis of FAs, membrane biogenesis and cell division regulation through histone degradation in protozoan parasites and mammalian cells. In all eukaryotic cellular species, membrane lipid synthesis from precursor FAs and bi-layer assembly is a prerequisite process before or a concomitant step with the commitment for nuclear division and cytokinesis. The role of the checkpoint is to instigate a controlled reversible cell cycle arrest, if required, to ascertain the level of macromolecular resources and DNA damage corrections before mitotic commitment. In malaria parasites, impaired FAs import and compromised membrane biogenesis led to the cessation of DNA replication and arrest of nuclei formation ^12, 24^ (Figure. 1B & 1C). The transcript level of cell cycle related genes e.g., tubulin, PCNA, DNA polymerase and DNA directed RNA polymerase were downregulated when FAs import was inhibited in *P. falciparum* 3D7 parasites (Figure. S7, Excel sheet. S5). Upon getting access to FAs through normal import, cell cycle marker genes were upregulated significantly to resume cell division (Figure. S7, Excel sheet. S6). The arrest of DNA replication and daughter nuclei formation resulted due to the degradation of histone proteins in the parasite nucleus by the evolutionary conserved bacterial ClpP/ClpR protein quality control machinery. ClpP and ClpR in Plasmodium showed a distinct nuclear localization signal (Figure. S8A). Multiple sequence alignment across cell type of prokaryotes and eukaryotes seemed to suggest a significant amino acid sequence conservation in ClpP, suggesting an important role probably through structural conservation (Figure. S8B). Phylogenetic tree analysis of ClpP across cellular species seemed to suggest a greater conservation rather than deviation (Figure. S8C & S8D). Inhibition of FAs import and subsequent impaired membrane biogenesis led to hyperphosphorylation of arginine residues in histones possibly by a NPD1951-AK protein with the plasmodb gene ID. PF3D7_1409200 (Figure. 6E & 6F). The inhibition of arginine hyperphosphorylation by arginine kinase antibody resulted in reduced histone degradation (Figure. 6E & 6F), which suggested the importance of arginine phosphorylation in order to degrade histones through ClpP/ClpR activity. The hyperphosphorylation of histones resulted into the down gradation of the quality of histones and as a result subjected for proteostasis or quality control measures by ClpP/ClpR machinery in the parasite nucleus. The hyperphosphorylation of arginine was achieved possibly by newly discovered arginine kinase, NPD1951-AK in the malaria parasites. The lipid and membrane biogenesis mediated checkpoint regulation through histone degradation before the commitment for DNA replication and mitotic nuclei formation has opened an entirely new front of cell cycle checkpoint regulation at the onset of schizogony in malaria parasites and in the G2 phase, but before mitotic commitment in mammalian cells (Figure. 7).

**Figure 7.**
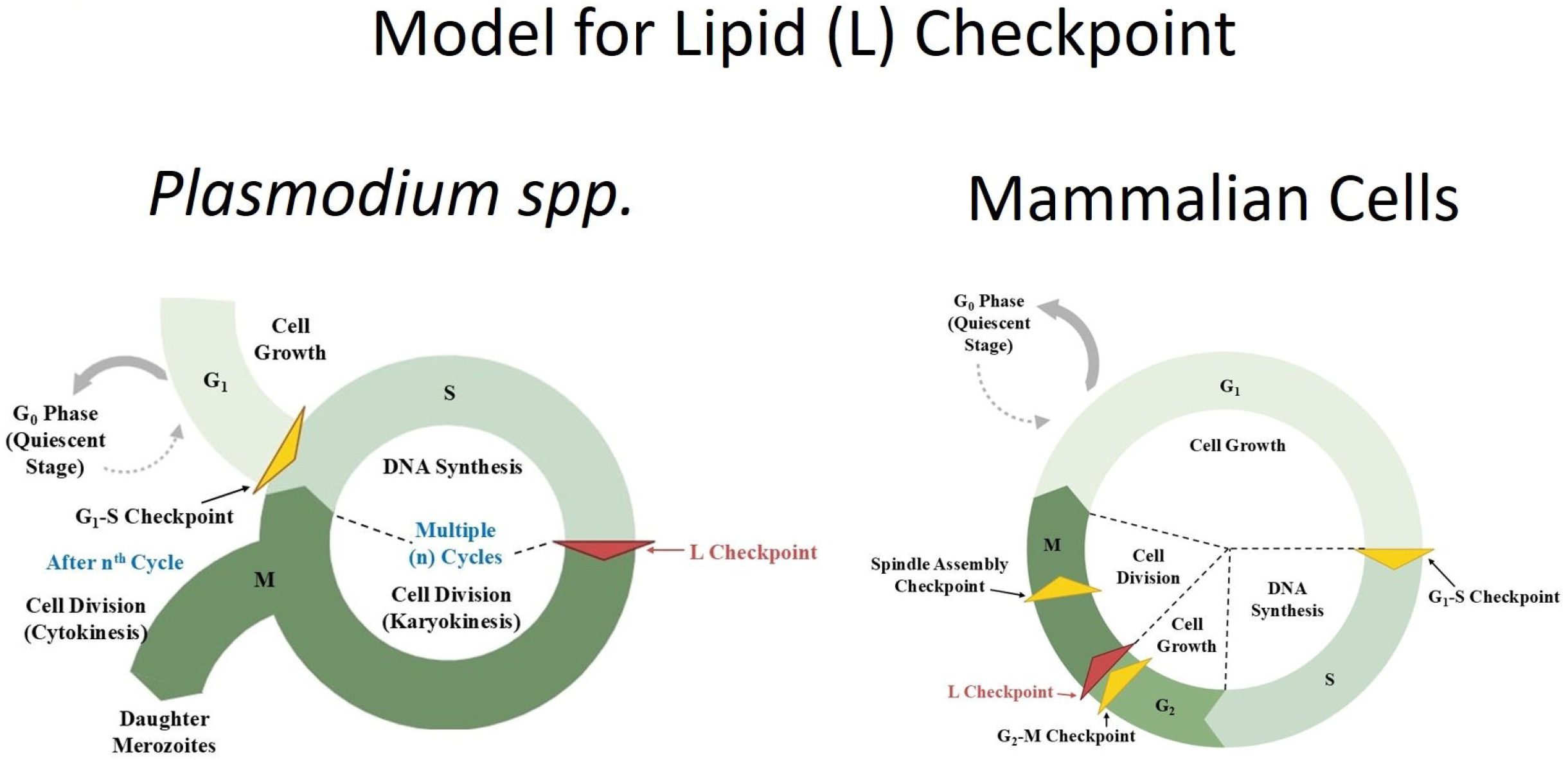
A novel cell cycle checkpoint in *Plasmodium spp* and mammalian cells coined as Lipid-checkpoint (L-checkpoint) **(A)** Model showing a cartoon of Cell cycle checkpoints in *Plasmodium spp* with an inclusion of our newly discovered Lipid (L) checkpoint. Between DNA synthesis (S phase) and nuclear division through Mitosis (M phase), lipid resources and homeostasis are checked for subsequent nuclear membrane assembly for daughter nuclei formation. Hence, the surveillance of lipid resources before the commitment to M phase are monitored in this newly proposed L-checkpoint. It appears that L-checkpoint decides whether after S phase cell should proceed towards M phase or go for temporary arrest before M phase until the lipid homeostasis is back to its normal. **(B)** Model showing cell cycle checkpoint in mammalian cells with an inclusion of our newly discovered Lipid (L) checkpoint after G2-M checkpoint. At S phase after DNA replication, before the commitment to M phase, L-checkpoint presumably make sure the availability of fatty acids and membrane lipids, which are required for daughter nuclear membrane formation after M phase and subsequent plasma membrane concomitant to cytokinesis. At L-Checkpoint, if there is lipid scarcity hence downregulation of Kennedy pathway for membrane lipid biosynthesis, cells appear to sense and signal for a pause until the lipids return back to its normal cellular level required to resume membrane biogenesis.

### Cell cycle checkpoints through lipid biogenesis in *P. falciparum*

Checkpoints are the opportunities to instigate a controlled arrest during the ongoing cell cycle process until the encountered problems are fixed. In malaria parasites, nuclei divide asynchronously, preceding cytokinesis and cellularization ^3, 4, 5^. Unlike mammalian cells, in malaria parasites, checkpoint mediated cell cycle regulation and the involvement of Cyclin-CDKs and cyclin related kinases (CRKs) are not well understood due to uncanny nature of cell cycle progression ^5, 6, 7, 9^. However, a recent study seems to suggest an indispensable role of CRK4 in the continuous round of DNA replication during S/M phase in the parasite ^64^. Checkpoint events in Plasmodium are very opaque and difficult to streamline. Recent evidences seem to suggest a controlled entry into the schizont stage as deprivation of isoleucine arrest cell cycle possibly at early schizont stage, which is reminiscent of an entry into the S phase via G1-S checkpoint ^65, 66^. However, recent evidences suggest that the arrest is not complete and the cells slowly proceed even under isoleucine deprived condition and initiate DNA replication ^66^. Polyamines, on the other hand, seem to play an important role in the parasite cell division as its deprivation arrest progression at the trophozoite stage ^67^. After repeated S/M phase, evidences seem to suggest that there may not be a cytokinesis checkpoint that would temporarily stall parasite cell cycle even if no viable daughter cells are formed ^67, 68, 69, 70^. Overall, in malaria parasites, there are no well accepted events which deemed fit to be called as checkpoints until recently some previous reports and our study showed that during IE parasite development, deprivation of FAs in the culture medium or the inhibition of FAs import into the iRBCs arrest parasite nuclear division at the transition of the trophozoite/schizont ^12, 24, 33, 34, 35, 36^. FAs deprivation through inhibition of import led to the accumulation of triacylglycerol (TAG) and consequentially downregulation of membrane lipid biosynthesis ^12^. The nature of crosstalk between nuclear division arrest and impaired membrane biogenesis via FAs deprivation was a most pressing unaddressed question which was hinting towards a checkpoint like event possibly at G1 phase which assures the level of lipid precursors for membrane bilayer synthesis of daughter nuclei before the commitment and/or during ongoing repeated nuclear division and mitosis (S/M phase). To explore how FAs deprivation and impaired membrane biogenesis led to the arrest of nuclear division, we undertook several approaches and established that the degradation of nuclear histones was a result of FAs deprivation and impaired membrane biogenesis. Evolutionary conserved parasite nuclear localized bacterial serine protease ClpP/ClpR degraded histones as a protein quality control measures upon finding hyperphosphorylated arginine residues in histones (Figure. 4 & 5 &6 & S6A). Newly discovered arginine kinase appears to localize in the parasite nucleus (Figure. 6C & 6D) and responsible for arginine hyperphosphorylation and marks histone for degradation by ClpP/ClpR machinery. Mechanistic understating as to how histone was degraded when FAs import was inhibited was comprehensively achieved but how arginine kinase was triggered to hyperphosphorylate histones under FAs deprived condition is not yet clear and, whether LncRNAs mediated transcription regulation of essential genes required to achieve that critical task is also yet to be unravelled.

### Cell cycle checkpoints and lipid homeostasis in mammalian cells

Massive membrane rearrangements and accumulation of lipids are the hallmark features of dividing mammalian cells^13^. Several membrane lipids rearrange during cell cycle progression for plasma membrane assembly as the perturbation of the membrane triggers a checkpoint event possibly at G1 phase ^13, 14^. G1, S and G2 checkpoints and its importance for the assessment of resources and DNA damage corrections in mammalian cells are well defined but whether problem in the membrane lipid resources and subsequently as a consequence the impaired membrane biogenesis has the ability to instigate a controlled cell cycle arrest was not known. Following our discovery in malaria parasites, we wanted to check the status of histones when *de novo* biosynthesis of FAs was inhibited using anticancer drug Cerulenin and another compound C75 in cultured HEK293T mammalian cells and carcinoma cell lines HCT116 to assess the effect of impaired membrane biogenesis on the cell cycle regulation and the quality of histone proteins (Figure. S1). To our surprize, inhibition of *de novo* FAs biosynthesis led to the degradation of histones and the arrest of cell proliferation (Figure. S1A & S1B). Interestingly, when FAs biosynthesis was impaired, HEK293T cell number was 3-4 times less and HCT116 cells were 5 times reduced after Cerulenin treatment (Figure. S1A & S1B). Paradoxically, mean fluorescence intensity (MFI) of nuclear stain DAPI was similar across all the samples, i.e., DMSO treated (control), Cerulenin and C75 treated (Figure. S1A & S1B). This observation was mindboggling hence raised a legitimate question about how significantly less numbers of cells showed a comparable MFI of nuclear stain DAPI. The probable explanation would be deeply hidden inside and we speculated that the inhibition of *de novo* FAs biosynthesis did not arrest DNA replication at “S” phase. But, at G2 phase, deprived FAs signalled and resulted into the arrest at “M” phase due to the inhibition of membrane lipid formation hence impaired membrane bi-layer assembly for daughter nuclei formation and plasma membrane synthesis. From this observation (Figure. S1A & S1B), it was evident that either histone degradation or impaired membrane biogenesis was equally capable of cell cycle arrest but it is equally likely that both conditions together resulted into cell division arrest. However, it is not yet clear, how replicated DNA assembled nucleosomes when histones were degraded or was it merely the duplication of genetic materials without segregation and spatio-temporal arrangement in the ongoing nuclei formation process. Histone degradation through the inhibition of *de novo* FAs biosynthesis in HEK293T and HCT116 cells raised a legitimate question whether the degradation was carried out by ubiquitin-Proteasome system. Immunoblot using anti-ubiquitin antibody of DMSO (control) and Cerulenin treated HEK293T cells seemed to suggest no difference in protein band pattern between these two conditions (Figure. S6B), hence it was likely that in the nucleus of mammalian cells the histone degradation was carried out by an unknown protease without the participation of ubiquitin-Proteasome system. If it was a protease then what was the identity? Interestingly, in the Cerulenin treated HEK293T cells, the expression level of ClpP and arginine kinase was reduced (Figure. S6B). With these mechanistic understating, we were inclined to speculate that we would not be surprized if the nucleus of a cell in all cellular species possess a proteostasis mechanism for nuclear proteins besides the proteosome-ubiquitin system.

Overall, we observed a remarkable dichotomy in the mechanism of cell cycle arrest through histone degradation, FAs deprivation and, impaired membrane biogenesis between Plasmodium and mammalian cells. Both the cell types were arrested but appeared to be at different stages due to two different reasons. In Plasmodium, arrest resulted into the cessation of DNA replication (Figure. 1B) possibly to inhibit repeated “S/M” phase and avoid the wastage of energy resources required for DNA replication and also avoid a polyploidy situation because there were no lipid components to assemble the nuclear membrane for daughter nuclei segregation. On the other hand, in mammalian cells, arrest most likely did not stop DNA replication as we observed similar MFI of nuclear stain DAPI (Figure. S1B), which seemed to suggest a similar number of duplication cycles of genetic material as in untreated cells. However, in the Cerulenin and C75 treated cells, cell number was significantly less (Figure. S1B), which further confirmed that the arrest was at the level of concomitant nuclear and plasma membrane assembly and cytokinesis.

### Lipid (L) checkpoint

Lipid resources for membrane biogenesis and their easy accessibility facilitate membrane bi-layer assembly during cellular proliferation. Nuclear membrane assembly, organellar membrane biogenesis, and plasma membrane biosynthesis, all are critical processes in the regulation of the cell cycle before the commitment for mitosis. Deprived FAs and impaired membrane lipid biosynthesis appeared to instigate a controlled cell cycle arrest either within the “G2” phase or between G2 and M phase, where the availability of lipid resources is strictly measured before the decision of full commitment for mitosis. We name this phase the Lipid (L) checkpoint (Figure. 7). In our study, we have implied the importance and the role of an L-checkpoint before mitotic commitment. Since FAs and membrane lipid biosynthesis are both fundamental processes, we believe that L-checkpoint is plausibly operational in all eukaryotic cellular species during cell cycle regulation prior to mitotic commitment.

### Limitation of the study

In this study, we have discovered a novel cell cycle checkpoint between G2 and M phase. We named this novel checkpoint the Lipid (L)-checkpoint, which is mediated by FAs and membrane biogenesis. Deprived FAs and impaired membrane biogenesis led to the arrest of cell cycle through histone degradation by evolutionary conserved bacterial protein quality control machinery ClpP/ClpR upon hyperphosphorylation of arginine residues in histones by a novel arginine kinase in the parasite nucleus. Although we have discovered the entire mechanism of histone degradation and cell cycle arrest, which was triggered by FAs deprivation, we still do not know the interplay between arginine kinase and a phosphatase, which was required for the regulation of phosphorylation and how arginine kinase is getting activated and transcriptionally regulated. In untreated parasites, when FAs and membrane biogenesis are normal, even in the presence of arginine kinase, how arginine residues remained protected from hyperphosphorylation, but upon FAs deprived condition, that protection seemed to disappear, and as a consequence, the arginine residues in histones were hyperphosphorylated. Although to address this issue, we have discovered the entire *P. falciparum* LncRNA-oms, but currently we do not have a direct correlation between LncRNA regulation and arginine kinase transcription. To add more, (1) structurally, we are not sure how hyperphosphorylated arginine residues in histones are playing a role in recognizing ClpP/ClpR machinery in the parasite nucleus for histone degradation. (2) In mammalian cells, ClpP does not localize in the nucleus, so how histones were degraded upon inhibition of *de novo* FAs biosynthesis. It opened an entirely new direction towards the histone quality control system in the cell nucleus, which was triggered by FAs deprivation and impaired membrane biogenesis.

## Figure Legends

**Supplementary Figure 1. Inhibition of *de novo* FAs biosynthesis led to histone degradation and arrest of cell division in mammalian cells HEK293T and human colorectal carcinoma cells HCT116, related to Figure 1 and 4**.

**(A)** Immunoblot of cultured mammalian cells HEK293T and HCT116 after DMSO (Control), Cerulenin (100µM) and C75 (350 µM) treatment. Blot was probed with anti-histone H2A antibody, anti-histone H2B antibody, anti-histone H3 antibody and anti-Histone H4 antibody. GAPDH was used as a loading control. **(B)** Flow cytometry analysis of DAPI stained nucleus of cultured mammalian cells HEK293T and HCT116 after DMSO (Control), C75 (350 µM) and Cerulenin (100µM) treatment. Total number of cells counted and mean fluorescence intensity (MFI) of DAPI staining cells was depicted. Gated single cells were used for DAPI count **(C)** Immunofluorescence assay (IFA) of ClpP protein in HEK293T cultured cells after DMSO, C75 (350 µM) and Cerulenin (100µM) treatment. ClpP was stained by anti-ClpP antibody and cell nucleus was stained with DAPI.

**Supplementary Figure 2. Induced gametocytogenesis through genetic knockdown of PfP2 protein in *P. falciparum* NF54 parasites, related to Figure. 1, 2 and 3**.

**(A)** Knockdown strategy of ribosomal protein P2 in *P. falciparum* NF54 parasites using pL-6 3xHA-glmS Ribozyme plasmid. **(B)** Diagnostic PCR to detect hDHFR and glmS Ribozyme integration in the *P. falciparum* NF54 genome and conformation of P2 gene in the integrated parasite line. **(C)** % parasitaemia count of *P. falciparum* NF54 parasites with or without 3 mM glucosamine (GlcN) treatment up to day 4. **(D)** Immunoblot of PfP2-3xHA glmS transgenic *P. falciparum* NF54 parasites with anti-HA and anti-PfP2 mAb E2G12 with or without GlcN treatment. Aldolase was used as a loading control. **(E)** Giemsa-stained transgenic *P. falciparum* NF54 parasite at PMI ~ 32-36h after P2 protein down regulation using 3 mM GlcN. Giemsa-stained parasites was a representative image.

**Supplementary Figure 3. Patatin-like phospholipase 1 (PATPL1) interacts with PK6 and PK9 and presumably regulates TAG level in the parasites, related to Figure 2**.

**(A)** Immunoblot of transgenic *P. falciparum* 3D7 Protein Kinase 6 (PK6-2xHA) and Protein Kinase 9 (PK9-2xHA). Blot was probed with anti-HA antibody. **(B)** Schematic showing the strategy of Co-Immunoprecipitation assay (Co-IP) followed by mass spectrometry ESI MS/MS of PK6-2xHA and PK9-2xHA using anti-HA antibody and Protein G Sepharose beads to detect interactors of PK6 and PK9. **(C & D)** Table showing mass spectrometry based identified interactor proteins of PK6 and PK9. **(E)** Heatmap of differentially expressed mRNAs after orlistat treatment in *P. falciparum* 3D7 parasites **(F)** Relative abundance of transcripts of XL1/XL2/PATPL1 and PATPL4 in untreated (control) and E2G12 treated *P. falciparum* 3D7 parasites. Statistical P values were denoted as *. P<0.05 as *, P<0.01 as **, P<0.001 as *** and P<0.0001 as ****.

**Supplementary Figure 4. Long-non coding RNAs (LncRNAs) respond to FAs import inhibition and presumably regulate gene expression according to the parasite physiology in *P. falciparum* 3D7, related to Figure 3**.

**(A)** Heatmap of 50 most differentially expressed *P. falciparum* 3D7 LncRNAs at trophozoite stage after E2G12 treatment. The scale of the LncRNA expression level was from 0 to 10, where 0 was comparable to control (untreated) and 10 depicts the highest expression of the LncRNA transcripts. **(B)** log2 fold change of 27 most differentially expressed LncRNA transcripts at trophozoite stage of *P. falciparum* 3D7 parasites showing significant upregulation and downregulation of LncRNAs after FAs import inhibition due to the blockage of PfP2 complex by E2G12 **(C & D)** Volcano and scatter plot of differentially expressed LncRNAs in *P. falciparum* 3D7 trophozoite parasite under untreated (control) and E2G12 treated conditions. **(E)** log2 fold change of LncRNA transcripts at trophozoite stage of *P. falciparum* 3D7 parasites after the rescue from E2G12 treatment showing significant upregulation and downregulation of LncRNAs. **(F)** log2 fold change of the same 27 most differentially expressed LncRNA transcripts as panel B at trophozoite stage of *P. falciparum* 3D7 parasites after the rescue from E2G12 treatment and restoration of FAs import. (G & H) Volcano and scatter plot of differentially expressed LncRNAs in P. falciparum 3D7 trophozoite parasite after the rescue from E2G12 treatment. (I & J) Table depicting chromosomal origin of differentially expressed LncRNAs in *P. falciparum* 3D7 parasites after the inhibition of fatty acid import by E2G12. Statistical P values were denoted as *. P<0.05 as *, P<0.01 as **, P<0.001 as *** and P<0.0001 as ****.

**Supplementary Figure 5. Differentially expressed LncRNAs in *P. falciparum* NF54 parasites and its identification after FAs import inhibition by E2G12, related to Figure 2**.

**(A)** Heatmap of 50 most differentially expressed *P. falciparum* NF54 LncRNA transcripts at trophozoite stage after E2G12 treatment. The scale of the LncRNA expression level was from 0 to 14, where 0 was comparable to control (untreated) and 14 depicts the highest expression of the LncRNA transcripts. **(B & C)** Volcano and scatter plot of differentially expressed LncRNAs in *P. falciparum* NF54 trophozoite stage parasites under untreated (control) and E2G12 treated conditions. **(D)** Log2 fold change of LncRNA transcripts showing significant upregulation and downregulation at trophozoite stage in *P. falciparum* NF54 parasites after FAs import inhibition by E2G12. Statistical P values were denoted as *. P<0.05 as *, P<0.01 as **, P<0.001 as *** and P<0.0001 as ****. **(E)** Table showing chromosomal origin of differentially expressed LncRNAs at trophozoite stage of *P. falciparum* NF54 parasites after the inhibition of FAs import by E2G12.

**Supplementary Figure 6. Evolutionary presence of ClpP protease, arginine kinase and creatine kinase enzymes in *E. coli, Cyanobacteria* and mammalian cells, related to Figure. 3, 4, 5 & 6**.

**(A)** Immunoblot of *E. coli* and *Cyanobacteria* lysate using Anti-ClpP, Anti-Arginine Kinase and Anti-Creatine Kinase antibody. **(B)** Immunoblot of HEK293T mammalian cells DMSO control Vs Cerulenin treated (100 µM) using anti-Ubiquitin antibody, anti-ClpP and anti-Arginine Kinase antibody. GPADH was used as a loading control. **(C)** IFA of *E. coli DH5*α cells using anti-ClpP antibody **(D)** IFA of *E. coli* cells using anti-Arginine Kinase antibody, raised against Arginine Kinase protein in Honeybee (*Apis mellifera*). **(E)** IFA of Cyanobacteria cells using anti-ClpP antibody **(F)** IFA of Cyanobacteria cells using anti-Arginine Kinase antibody **(G)** IFA of *E. coli* cells using anti-Creatine Kinase antibody (H) IFA of Cyanobacteria cells using anti-Creatine Kinase antibody.

**Supplementary Figure 7. Gene expression and transcripts abundance of cell cycle related genes in *P. falciparum* 3D7 trophozoite stage parasites after FAs import inhibition and rescue, related to Figure. 1**.

**(A)** Relative abundance of transcripts of cell cycle related genes after FAs import inhibition using anti-PfP2 mAb E2G12 at trophozoite stage of *P. falciparum* 3D7. **(B)** Relative abundance of transcripts of cell cycle related genes at trophozoite stage of P. falciparum 3D7 parasites after the rescue from E2G12 treatment. Statistical P values were denoted as *. P<0.05 as *, P<0.01 as **, P<0.001 as *** and P<0.0001 as ****.

**Supplementary Figure 8. Multiple sequence alignment and phylogenetic tree analysis of ClpP and arginine kinase across different cell types and identification of nuclear localization signal in *P. falciparum* 3D7, related to Figure 4, 5, & 6**.

**(A)** Nuclear localization signal of ClpP (43 kDa) and ClpR (28 kDa) in *P. falciparum* 3D7 parasites. **(B)** Multiple sequence alignment of ClpP across prokaryotes, in humans and in *P. falciparum* showing amino acid sequence similarities in ClpP protein (red) and evolutionary conservation. **(C)** Phylogenetic tree of ClpP protein showing relatedness across bacteria, in humans and in *P. falciparum* **(D)** Phylogenetic tree analysis of ClpP across cell types. The relatedness across clades and species within a clade were depicted by bootstrap values signifying how close ClpP protein in different organism. **(E)** ClustalW of Arginine Kinase in Honeybee (*Apis mellifera*) and in other cell types showing amino acid sequence similarity and identity **(F)** Phylogenetic tree analysis of Arginine Kinase with bootstrap values and relatedness across different cell types. **(G)** Amino acid sequence alignment of adenylate kinase isoform 6 in human, adenylate kinase like protein and conserved proteins in *P. falciparum* **(H)** Predicted nuclear localization signal (blue) in human adenylate kinase isoform 6 adenylate kinase like protein and conserved proteins in *P. falciparum*. Nuclear localization signal was predicted using NLS mapper software (https://nls-mapper.iab.keio.ac.jp/cgi-bin/NLS_Mapper_form.cgi).

## Supplementary Excel Data Sheets

### Excel Sheet. S1

E2G12 dose dependent treatment of *P. falciparum* 3D7 parasites for LC-MS lipid profiling at each antibody concentration and identification of lipid species.

### Excel Sheet. S2

Transcriptomics profiling of Untreated (Control) Vs E2G12 treated (FAs import inhibited) *P. falciparum* NF54 trophozoite stage parasites with log2 fold upregulation/ downregulation of 4745 genes / transcripts including apicoplast and Mitochondria genes and identification of each gene.

### Excel Sheet. S3

Transcriptomics profiling of Untreated (Control) Vs Orlistat treated *P. falciparum* NF54 trophozoite stage parasites with log2 fold upregulation/ downregulation of 4538 genes / transcripts including apicoplast and Mitochondria genes and identification of each gene.

### Excel Sheet. S4

E2G12 dose dependent treatment of *P. falciparum* NF54 parasites for LC-MS lipid profiling at each antibody concentration and identification of lipid species.

### Excel Sheet. S5

Transcriptomics profiling of Untreated (Control) Vs E2G12 treated (FAs import inhibited) *P. falciparum* 3D7 trophozoite stage parasites with log2 fold upregulation/ downregulation of 4546 genes / transcripts including apicoplast genes and identification of each gene.

### Excel Sheet. S6

Transcriptomics profiling of E2G12 treated (FAs import inhibited) Vs E2G12 Rescued *P. falciparum* 3D7 trophozoite stage parasites with log2 fold upregulation/ downregulation of 5272 genes / transcripts including apicoplast and Mitochondria genes and identification of each gene.

### Excel Sheet. S7

Transcriptomics profiling of Untreated (Control) Vs Orlistat treated *P. falciparum* 3D7 trophozoite stage parasites with log2 fold upregulation/ downregulation of 4573 genes / transcripts including apicoplast and Mitochondria genes and identification of each gene.

### Excel Sheet. S8

qTOF ESI-MS mass spectrometry-based profiling of Protein Kinase 6 (PK6) and PK9 interactors proteins and their identifications.

### Excel Sheet. S10

Transcriptomics profiling of LncRNAs in *P. falciparum* 3D7 at trophozoite stage parasites after Control Vs E2G12 treatment with log2 fold upregulation / downregulation of transcripts expression.

### Excel Sheet. S11

Transcriptomics profiling of LncRNAs in *P. falciparum* 3D7 at trophozoite stage parasites after E2G12 treatment Vs E2G12 Rescued with log2 fold upregulation / downregulation of transcripts expression.

### Excel Sheet. S12

Transcriptomics profiling of LncRNAs in *P. falciparum* NF54 at trophozoite stage parasites after Control Vs E2G12 treatment with log2 fold upregulation / downregulation of transcripts expression.

### Excel Sheet. S13

qTOF ESI MS/MS mass spectrometry-based Post Translational Modification (PTMs) profiling of Ac Et.OH extracted Histone H2A, H2B, H3 and H4 of Untreated and E2G12 treated *P. falciparum* 3D7 trophozoite stage parasites.

### Excel Sheet. S14

qTOF ESI-MS mass spectrometry-based profiling to identify Arginine Kinase and its interactor proteins in *P. falciparum* 3D7 trophozoite stage parasites.

## Materials and Methods

### Generation of anti-PfP2 mAb E2G12 and rabbit polyclonal antibodies

Anti-PfP2 mAb E2G12 (Das et al., 2012a, 2012b, 2024)^12, 25, 26^ was custom generated by Bioklone Biotech India Pvt. Company, based in Chennai, India.

### *P. falciparum* 3D7 and NF54 parasite culture

*P. falciparum* 3D7 and NF54 parasites were cultured using human RBCs (type O^+^) in RPMI1640 media supplemented with sodium bicarbonate (2 g/L), glucose (2g/L), hypoxanthine (10 mg/L), gentamicin sulphate (50 µg/L), and 0.5% Albumax (cRPMI). Asexual stages of *P. falciparum* 3D7 and NF54 parasites were cultured using RBCs 5% haematocrit in cRPMI and maintained in a 37°C humidified incubator under 89% N2, 5% O2 and 6% CO2. Periodic testing for Mycoplasma contamination by PCR was used to ensure that growing parasite cultures were free from such contaminations. Growing parasite cultures were synchronized at the ring stage using 500 mM Alanine, 10 mM HEPES, pH = 7.4. Studies were conducted using ring stage parasites at 6–7% parasitaemia following two rounds of synchronization. The *P. falciparum* Protein Kinase (PK6) and PK9 transgenic 3D7 parasite lines were propagated in RPMI media containing Blasticidin-HCl (5 µg/mL) under culture conditions as described above.

### Culture of mammalian cell HCT116 and HEK 293T and treatment with C75 and Cerulenin

Both HCT116 and HEK293T cell lines were grown in a petri dish in a CO_2_ incubator under humidified condition. 5% CO_2_ was maintained for optimum growth condition. To grow cells, DMEM was used as a media and cells were passaged after every 48-72h. Cells were harvested after trypsinization followed by biochemical experimentations. HEK293T and HCT116 cells were cultured in DMEM and equal number of cells were treated with DMSO control, 350 uM C75 and 100 uM Cerulenin. Both the compounds were made soluble in pure DMSO. After 24h of treatment at 37°C in the CO2 incubator, cells were harvested after trypsinization and centrifugation. The cell pellets were subsequently washed once with ice cold 1x PBS and pelleted down. The final cell pellets were lysed though sonication in 1x PBS with/without 0.1% Triton X100 and with/without protease inhibitor. The cell lysates were centrifuged at 14000 *xg* at 4°C for 30 mins and supernatant was collected. Protein was estimated using BCA and quantified. Equal amount of protein was separated in SDS-PAGE and western blot was performed using histone specific antibodies H2A, H2B, H3 and H4. As a loading control GAPDH was used.

### Plasmid construction for DNA electroporation

For conditional knockdown of the PfP2 protein in *P. falciparum* 3D7^12^ and NF54, homology region 1 (HR1) and HR2 were PCR amplified from the coding and intergenic region and the 3’UTR respectively with two sets of primers. Both the homology regions were cloned into the pL6-3HA-glms-ribozyme vector which contains human dihydrofolate reductase (hDHFR) as a selectable marker that confers resistance to WR99210. HR1 and HR2 PCR products were cloned one by one with the sequencing of DNA. A list of prospective 20 base nucleotide sequences (N20) for guide RNAs was generated using the Eukaryotic Pathogen CRISPR guide RNA (gRNA) design tool (http://grna.ctegd.uga.edu/) that targets the *P2 gene* in the chromosomal DNA segment flanked by the 5’ and 3’ HRs. gRNAs were cloned in pUF1-Cas9 vector which carries a gRNA expression cassette, Cas9 endonuclease expression cassette and a yeast dihydroorotate dehydrogenase (yDHODH) cassette for selection with DSM1. For the oligonucleotide sequences used to generate plasmid constructs were mentioned ion supplementary table 1.

### Parasite line, transfection method, transgenic parasite line selection and PfP2 downregulation

*P. falciparum* 3D7^12^ and NF54 parasites were used to generate transgenic parasites. Standard transfection method, parasite selection and negative selection with 5FC and limiting dilution to select integrated parasites were followed as described in Prommana et al., ^74^ and Ito et al., ^75^. Briefly, ring-stage parasites from a culture at 6-7% parasitaemia were washed three times with pre-warmed Cytomix (pH =7.6) and then resuspended with an equal volume of ice-cold Cytomix. An aliquot of 250 µl of ring stage parasite suspension was mixed with 50 µg of both the plasmids and put in 0.2 cm cuvette for electroporation. BioRad Gene Pulser was set at 0.31 kV and 950 μF. Parasite cultures were selected using 5 nM WR99210 and 1.5µM DSM1. After 2-3 weeks of transfection, parasites were detected using Giemsa staining. PCR was used to evaluate the integration of the hDHFR and glmS. For Conditional downregulation of P2 protein under the *glmS* ribozyme system, 3 mM GlcN was added to synchronous ring-stage parasites at around 12-14h PMI (Prommana et al.,^74^). GlcN exposure was continued for up to 10h before washout and rescue experiments or kept continued before harvest for phenotype studies fixation and IFA and biochemical experiments. Control experiments with GlcN added in the culture media revealed no measurable toxicity in wild-type *P. falciparum* 3D7 parasites at up to a 4 mM concentration.

### Treatment of *P. falciparum* 3D7 and NF54 infected RBCs with mitotic inhibitor or cell cycle arrester for Growth Inhibition assay (GIA)

Anti-PfP2 mAb E2G12 was salted out using 50% ammonium sulphate solution and kept overnight for complete precipitation at 4°C. The solution was centrifuged at 10,000 *xg* at 4°C. The precipitated antibody E2G12 was then resuspended in 1x PBS and dialyzed against 1x PBS to remove ammonium sulphate. Dialysed antibody was quantified using BCA kit. *P. falciparum* 3D7 and NF54 parasites were cultured till 6-8% parasitaemia and synchronized two rounds using 500 mM Alanine and 10 mM HEPES pH 7.4 as previously mentioned (Das et al., 2024)^12^. Synchronized ring stage parasites after two generations were treated with E2G12 (700 ng/µL) or Orlistat (20 µM) for 24h (arrested condition) or treated for 24h followed by removal of antibody through washing with normal media and re-cultured in antibody free media for another 12h (rescue condition). Following treatment, the iRBC pellets were collected, washed with 1xPBS and were resuspended in 1x PBS containing 0.1% saponin with added protease inhibitor cocktail (PIC) followed by 10 min incubation at 37°C to allow RBC lysis. The lysates were then centrifuged for 15 min at 10,000*xg* at RT (room temp.) to collect released parasites. The parasite pellet was then washed twice with 1x PBS. These pelleted parasites were later processed for western blotting or kept in −80°C for future usage. Similarly, *P. falciparum* 3D7 and NF54 parasite infected RBCs were treated with a variety of mitotic inhibitors or cell cycle arrester for 24h at 37°C. The different treatments include Orlistat (20 µM), Aphidicolin (2 µg/ml) and Taxol (500 nM). As mentioned, iRBCs were processed with 0.1% saponin in 1x PBS and parasites were harvested.

### Treatment of *P. falciparum* 3D7 and NF54 infected RBCs with anti-PfP2 mAb E2G12 for time point assessments

Doubly synchronized *P. falciparum* 3D7 and NF54 parasites at 10-14h PMI at ring stage were treated for varying time with E2G12 (700 ng/ul) under GIA condition. E2G12 treatment was started at 10-14h PMI which was considered as 0h and thereafter parasites were harvested after every 6h till 36h. Collected iRBCs were treated with 0.1% saponin in 1x PBS and parasites were harvested after washing with 1x PBS twice. Harvested parasites were subsequently used for several biochemical experiments.

### Western blotting

The protein samples were resolved in 10% SDS-PAGE. Proteins were then transferred on to methanol-activated polyvinylidene fluoride (PVDF) membrane for 2h at 4°C via wet transfer method using chilled transfer buffer (25 mM Tris, 192 mM glycine,10% Methanol). The membranes were blocked with 5% BSA in 1xPBS overnight and then incubated on a rocker with primary antibodies with required dilution in 1xPBS containing 0.2% Tween 20 (PBST) for 3 h at room temperature (RT). Membranes were washed with PBST and then incubated with appropriate horseradish peroxidase (HRP) conjugated secondary antibodies (1:10,000 in PBST). Membranes were washed with PBST for 5 min at least 5–6 times and the blots were developed using the chemiluminescent substrate. Antibodies used were: anti-PfP2 E2G12 mAb: 1:1K; anti-PfAldolase antibody: 1:5K; anti-β-actin: 1:2K; anti-β-tubulin: 1:2K; anti-Histone H2A: 1:5K, anti-Histone H2B: 1:5K, anti-Histone H3, 1:5K; Histone H4: 1:5K; Anti-HA tag antibodies 1:1K. Blots were imaged using ChemiDoc system from Invitrogen.

### Cross-Incubation Lysate-Lysate degradation (CILLD)

*P. falciparum* 3D7 parasite pellets of untreated (control), Orlistat treated and E2G12 treated were sonicated as mentioned previously and then centrifuged at 12000*xg* at 4°C. The lysates were prepared without the addition of protease inhibitor cocktail (PIC). These lysates were inter-mixed in the following combinations – 5µL of control lysate + 10 µL E2G12 treated lysates (C+E), 5 µL control lysate + 10 µL Orlistat treated lysates (C+O), and 5µL of control lysate + 10 µL E2G12 treated lysates with PIC. As a control, 5 µL control lysate + 10 µL Buffer was mixed. In all the combinations, total amount of protein was same. These inter-mixed lysates were then incubated for 6h at 37°C. After treatment, the lysate combinations were analysed via western blotting using anti-Histone H3, anti-Histone H4 antibodies to check any degradation of histones.

### Lipidomic analysis by GC MS / LC MS

Synchronized *P. falciparum* 3D7 and NF54 parasite iRBCs were treated with E2G12 at a concentration of 700 ng/µL for 24 h. Untreated parasites were considered as control. After treatment, parasites were harvested with 0.1% saponin in 1x PBS and centrifuged to get the parasite pellet. Thereafter, the pellet was subjected for GC-MS and LC-MS analysis for the detection of fatty acids and lipids as described in Hara and Radin^76^. Parasite pellets were processed by adding a mixture of 2 mL Isopropanol: Hexane (1: 2)and 2ml of KCl (2M): MeOH at a ratio of 4:1. Samples were mixed and centrifuged at 4000 rpm for 5 mins. The organic top layer was collected in a chromatography vial and the pellet was re-extracted 3 times in 2 ml isopropanol: hexane 1:2. The organic phase was dried with nitrogen gas and resuspended in 500 µl hexane. The samples were dried again with nitrogen gas and resuspended in 300 µL chloroform: methanol (2: 1). Lipid extracts were analyzed using a Triple Quadrupole LC/MS system using electronspray ionization in the negative ion mode and were spiked with appropriate internal standards. In GC MS analysis, a Shimadzu USA GC MS QP 2020 model was used which is equipped with a SHRTx5 Shimadzu column(30m X 0.25mm X 0.25 µm). At different retention times, peaks were observed in GC-MS and LC MS corroborated with adduct masses of fatty acids (FAs) and phospholipids. In another set up, NF54 parasite infected RBCs at the Post Merozoite Invasion (PMI) stage were treated with anti PfP2 mAb (700 ng/µL) at different time points (0h, 6h, 12h, 24h, 30h,36h) and parasite pellets were subjected to saponin lysis. Furthermore, Giemsa staining of the blood smears were done where gametocyte formation at different time points were visualized after antibody treatment and lipid profile of each sample were assessed.

### Nano-LC-ESI MS/MS

IP eluates from iRBC membrane proteins, parasite membrane proteins or parasite cytosol were resolved in a 10% SDS PAGE or a 7% Native PAGE. Gels were stained with freshly prepared Coomassie Blue R250 stain or silver staining was performed as required. Bands were carefully excised and Coomassie stained gel pieces were consecutively washed with 100 mM Ammonium bicarbonate at 37°C for 15 min followed by 50% acetonitrile in 100 mM Ammonium bicarbonate for 15 min at 37°C. Gel pieces were washed in this manner until complete destaining. Silver stained gel pieces were incubated with destainer A and B following the manufacturer’s protocols followed by multiple washes with ultrapure water for 5 min. Destained gels were incubated with 20 mM DTT in 100 mM Ammonium bicarbonate and incubated at 60°C for 30 min for reduction. These gel pieces were then incubated with 50 mM Iodoacetamide (IAA) in 100 mM Ammonium bicarbonate at room temperature for 45 min in the dark for alkylation. Gel pieces were washed with 100 mM Ammonium bicarbonate and twice with 50% Acetonitrile in 100 mM Ammonium bicarbonate. Finally, gel pieces were completely shrunk by the addition of Acetonitrile before being dried in a laminar airflow hood. Completely dried gels pieces were rehydrated by the addition of Trypsin (20ng/ml) in 100 mM Ammonium bicarbonate and then incubated at 37°C for 10h. Precautions were taken to prevent any keratin contamination during the handling and the processing of the samples.

After trypsinization, the gel pieces were centrifuged to collect the supernatant which contained the tryptic digest. Peptides were purified using C18 spin column. Thereafter, peptides were extracted twice with 0.1% Formic acid in 70% Acetonitrile.

In some cases, in solution digestion of IP samples have been performed. Beads attached with IP proteins were directly incubated with 25 µL of DTT in Ammonium bicarbonate (reduction) followed by 35 µL of IAA in Ammonium bicarbonate (alkylation) followed by the addition of trypsin (trypsinization). The resulting solution contained trypsin digested peptides. Tryptic digests were collected from beads and was desalted using C18 spin columns. The C18 resin was activated using 200 µL of Acetonitrile containing 0.1% Formic acid, followed by centrifugation at 2300*xg* for 1 min and the process repeated. The collected tryptic digests were similarly passed through the columns followed by centrifugation and the step was repeated four times to ensure maximum peptide binding to the resin. These resins were then twice washed with 0.1% formic acid. Peptides bound to the resin were then eluted using 70% acetonitrile containing 0.1% formic acid. Elution was repeated again to ensure maximum peptide recovery. The MS/MS analyses of these C18 purified peptides were performed using ESI MS/MS in LTQ ORBITRAP XL Thermo Scientific or by qTOF, Waters system. Spectra were analyzed using Proteome Discoverer software version: 1.4.0.288 (Thermo). Searches were made using SequestHT and limited to the *P. falciparum* proteins sourced from UniProtKB.

### Total transcriptomics and RNA sequencing

*P. falciparum* 3D7 and NF54 parasites were cultured in human O^+^ RBCs and synchronized twice at the ring stage using 500 mM Alanine and 10 mM HEPES pH 7.4 solution as previously mentioned ^12^. Blood smears were prepared and Giemsa stained to ensure that cultures were at a high parasitaemia (7-8%) and predominantly at ring stage. The12-14h PMI ring stage iRBCs were treated for 24h under the following conditions –untreated (control), E2G12 700ng/µL) and Orlistat (20µM). In case of rescue experiments, iRBCs were extensively washed with media and re-incubated in E2G12 free fresh media and in new flasks for further 12h before being harvested. All treated and untreated (control) iRBCs were finally subjected to 0.1% saponin lysis in 1x PBS and parasite pellet were collected and stored at −80°C.**Total RNA extraction, Qualitative and Quantitative analysis:** Total RNA was extracted from parasite pellet using Direct-Zol RNA miniprep Kit (Zymo Research) as per manufacturer’s instruction. The quality and quantity of the extracted RNA were checked on NanoDrop followed by Agilent Tape station using High sensitivity RNA screen Tape. **Illumina PE Library preparation:** The paired end sequencing libraries were prepared from RNA samples using NEBNext rRNA depletion Kit and NEB Next Ultra ™ II Directional RNA library prep kit for Illumina (NEB)as per manufacturer’s instruction. Briefly, approximately, 1 ug of total RNA was processed for ribodepletion using NEBNext rRNA depletion kit, followed by enzymatic fragmentation, 1^st^ strand cDNA conversion using Super Script II and Act-D mix to facilitate RNA dependent synthesis as per the Illumina TruSeq Total RNA library Prep Kit. The 1^st^ strand cDNA wax then synthesised to second strand using second strand mix. The dscDNA was then purified using AMPureXP beads followed by A-tailing, adaptor ligation and then enriched by limited number of PCR cycles. **Quantity and Quality Check (QC) of Library on Agilent 4200 Tape station:** The PCR enriched libraries were purified using AMPureXP beads and analysed on 4200 Tape station system (Agilent Technologies) using high sensitivity D1000 screen tape as per manufacturer instructions. **Cluster Generation and Sequencing on Illumina platform:** After obtaining the Qubit concentration for the libraries and mean peak sizes from the Agilent Tape station profile, the PE Illumina libraries was loaded onto Illumina NovaSeq X Plus platform for cluster generation and sequencing. Paired-end sequencing allows the template fragments to be sequenced in both the forward and reverse directions on NovaSeq X plus. The kit reagents were used in binding of samples to complementary adaptor oligos on paired-end flow cell. The adaptors were designed to allow selective cleavage of the forward strands after resynthesis of the reverse strand during sequencing. The copied reverse strand was used to sequence from the opposite end of the fragment.

### ClpP and Histone dual Immunofluorescent assay (IFA)

*P. falciparum* 3D7 and NF54 parasites were cultured using complete RPMI and gas condition. Two round synchronized ring stage parasites at 10-14h PMI were treated with anti-PfP2 mAb E2G12 (700 ng/ul) for 24 h. After the treatment, untreated and E2G12 treated parasites were fixed using 4% paraformaldehyde and 0.0075% glutaraldehyde for 1 h at room temperature (RT). Fixed iRBCs were then permeabilized with 0.5% Triton X100 in 1X PBS for 1h at RT. Subsequent to that, permeabilized iRBCs were then blocked by 5% BSA in 0.1% Triton X100 containing IXPBS for 2h at RT. Blocked iRBC were then washed twice with 1X PBS and incubated with anti-ClpP antibody in 0.1% TritonX100 containing 1x PBS for 4h. After primary antibody incubation, iRBCs were washed twice with 0.1% Triton X100 containing 1x PBS. Thereafter, appropriate Alexa Fluor-488 conjugated secondary antibody was incubated in 0.1% TritonX100 containing 1x PBS for 2h. After secondary antibody incubation, iRBCs were washed twice with 1x PBS. To stain Histone 4 (H4), anti-mouse H4 antibody in 0.1% TritonX100 containing 1x PBS was incubated for 4h at RT. Subsequent to that, iRBCs were washed twice with 0.1% Triton X100 containing 1x PBS and then appropriate Alexa Fluor-647 conjugated secondary antibody was incubated in 0.1% Triton X100 containing 1x PBS for 2h. After that, iRBCs were washed twice with 1x PBS and incubated with DAPI (0.1 ug/ml) for 20 mins. Stained iRBCs were then finally washed twice with 0.1 % Triton X100 containing 1x PBS. Final parasite pellet was resuspended in 1x PBS for confocal imaging. iRBCs were imaged using a Leica confocal microscope. Model: Leica TCS SP8, Objective: 100×/60X, NA:1.4, PMT detector. Acquired IFA images were processed using LAS X and ImageJ software.

### Arginine Kinase Immunofluorescence Assay (IFA) in *P. falciparum* 3D7 and NF54 parasites

*P. falciparum* 3D7 and NF54 parasites were cultured using complete RPMI and gas condition. Two rounds synchronized trophozoite stage parasites were fixed using 4% paraformaldehyde and 0.0075% glutaraldehyde for 1 h at room temperature (RT). Fixed iRBCs were then permeabilized with 0.5% Triton X100 in 1X PBS for 2h at RT. Subsequent to that, permeabilized iRBCs were then blocked by 5% BSA in 0.1% Triton X100 containing 1x PBS for 2h at RT. Blocked iRBC were then washed twice with 1x PBS and incubated with anti-Arginine Kinase antibody (1:250) in 0.1% TritonX100 containing 1x PBS for overnight. After primary antibody incubation, iRBCs were washed twice with 0.1% Triton X100 containing 1x PBS. Thereafter, appropriate Alexa Fluor-647 conjugated secondary antibody was incubated in 0.1% TritonX100 containing 1x PBS for 2h. After secondary antibody incubation, iRBCs incubated with DAPI (0.1 ug/ml) for 20 mins. Stained iRBCs were then finally washed twice with 0.1 % Triton X100 containing 1x PBS. Final parasite pellet was resuspended in 1x PBS for confocal imaging. iRBCs were imaged using a Leica confocal microscope. Model: Leica TCS SP8, Objective: 100×/60X, NA:1.4, PMT detector. Acquired IFA images were processed using LAS X and ImageJ software.

### Immunoprecipitation (IP) using Arginine Kinase (AK) antibody and qTOF ESI-MS/MS to identify AK peptides and its interactors

After two rounds of synchronization, *P. falciparum* 3D7 ring stage parasites at 12-14h PMI were treated with E2G12 (700 ng/µl) antibody for 24h at 37°C under GIA condition. After the treatment, arrested parasites, were harvested using 0.1% saponin and parasite pellet was collected and washed twice with 1x PBS and pelleted by centrifugation. Parasite pellet was resuspended in lysis buffer containing 1x PBS having 0.1% Triton X100 and protease inhibitor cocktail. Parasite cells were sonicated and thereafter centrifuged at 14,000*xg* at 4°C. Supernatant was collected and total protein was quantified using BCA. Total 2 mg parasite protein was incubated with 4 µg of Arginine Kinase antibody at RT for overnight. On the other hand, Protein A Sepharose beads was washed with 1x PBS and blocked using 5% BSA in 1x PBS. Thereafter, beads were incubated with the parasite cytosol-arginine kinase antibody mixture for 4h at RT. Antibody bound Protein A beads were pelleted by centrifugation at 700*xg* at RT and washed 3-4 times with 1x PBS. The beads and antibody bound Arginine Kinase and its interactor proteins were directly digested using trypsin (20ng/ml) and thereafter peptides were collected and processed for ESI MS//MS as described under mass spectrometry method.

### Histone enrichment / isolation using acidified ethanol (Ac. EtOH) and qTOF MS/MS mass spectrometry for PTMs identifications

As before, synchronized *P. falciparum* 3D7 parasites maintained at high parasitaemia (7-8%) in T75 flasks were treated with anti-PfP2 mAb E2G12 (700 ng/µL), Orlistat (20 µM) or kept untreated (control). After 24h post treatment, the iRBCs from these flasks were harvested, and then subjected to saponin lysis. Following centrifugation, the released parasite pellets were washed with PBS and then stored at −80°C for further processing. As described in Homsi et al.,^55^, briefly, the parasite pellet was lysed in acidified ethanol through repeated vortexing and incubation in ice for 30 mins. The lysate was centrifuged at 4°C and supernatant was collected which predominantly had histones and other basic pI proteins and some metabolites. A 3 kDa PES membrane cut off filtration system was activated with buffer (Tris 20 mM and NaCl 100 mM) pH 7.4. The collected histones in acidified ethanol were enriched and purified using the 3 kDa cut off filter and gradually acidified ethanol was replaced by the buffer system. After 3-4 times of buffer exchange and histone enrichment, the collected histones in Tris-NaCl were speed vac and dried completely. The powder was kept at −80 °C for further mass spectrometry-based processing.

For ESI MS/MS processing, the enriched histones were re-dissolved in Ammonium bicarbonate buffer. Chymotrypsin at 50 nM concentration was used for peptide fragmentation overnight at 37 °C. The digested peptides were enriched and salt free using C18 spin column. Resin bound peptides were eluted in 70% ACN/Formic and loaded in nano-LC for mass spectrometry analysis using qTOF. Acquired data was analysed using Progenesis software from Waters and *Plasmodium falciparum* Uniprot database was used.

### GIA using E2G12 and CLPP expression analysis using anti-ClpP antibody in Western blots

*P. falciparum* 3D7 and NF54 parasites were cultured using complete RPMI and gas condition. Two rounds synchronized 10-14h PMI ring stage parasites were treated with anti-PfP2 mAb E2G12 (700 ng/ul) for 24h. Total number of parasites were kept same in control (untreated) and in E2G12 treated culture flasks. After the 24h of treatment, *P. falciparum* 3D7 parasites, were harvested using 0.1% saponin in 1x PBS. On the other hand, *P. falciparum* NF54 parasites were kept with E2G12 for additional 24h and altogether after 48h, parasites were harvested using 0.1% saponin in 1x PBS. Harvested parasites were washed twice with 1x PBS and the parasite pellet were lysed and SDS-PAGE was run and was subsequently western blot was performed using anti-ClpP antibody. Appropriate anti-rabbit secondary antibody was used for image development.

### Immunofluorescence Assay (IFA) of *E. coli* and *Cyanobacteria* using Anti-ClpP, Arginine Kinase and Creatine Kinase polyclonal antibody

Lab strain *E. coli* was grown in LB medium and bacterial cells were harvested by centrifugation. Bacteria pellet was resuspended in a fixing solution (4% Paraformaldehyde and 1% Glutaraldehyde) for 3h. Fixed bacterial cells were washed twice with 1x PBS and permeabilized with 0.5% TritonX100 in 1x PBS for 2h at RT. Permeabilized bacterial cell population was then blocked with 5% BSA in 0.1% TritonX100 containing 1xPBS for 2h at RT. Cell were then washed twice with 0.1% TritonX100 containing 1xPBS and then incubated with Anti-ClpP antibody (1:100), anti-Arginine Kinase antibody (1:300) and anti-creatine kinase antibody (1:500) in 0.1% TritonX100 containing 1xPBS for 3h at RT. After ClpP antibody incubation, cells were washed twice with 0.1% TritonX100 containing 1xPBS and then incubated with anti-rabbit Alexa-488 conjugated IgG in 0.1% TritonX100 containing 1xPBS for 2h at RT. Finally, bacterial cells were washed twice with 0.1% TritonX100 containing 1xPBS and then twice with only 1xPBS and resuspended in 1x PBS. Entire staining procedure was done in solution.

Cultured Cyanobacterial cells were similarly stained with Anti-ClpP and Anti-Arginine Kinase antibody. Cyanobacteria cells were fixed with a fixing solution (4% Paraformaldehyde and 1% Glutaraldehyde) for 3h. Fixed bacterial cells were washed twice with 1x PBS and permeabilized with 1% TritonX100 in 1x PBS for 2h at RT. Permeabilized bacterial cell population was then blocked with 5% BSA in 0.1% TritonX100 containing 1xPBS for 2h at RT. Cell were then washed twice with 0.1% TritonX100 containing 1xPBS and then incubated with Anti-ClpP, Anti-Arginine Kinase antibody and anti-creatine kinase antibody. Appropriate Anti-rabbit Alexa 488 conjugated secondary antibody was used for 2h at RT. Thereafter, cells were washed twice with 1x PBS and resuspended in 1x PBS for imaging using Leica confocal microscope. Model: Leica TCS SP8, Objective: 100×/60X, NA:1.4, PMT detector. Acquired IFA images were processed using ImageJ software and LAS X software.

### Flow Cytometry

*P. falciparum* 3D7 parasites were synchronized to achieve 12-14h Post Merozoite Invasion (PMI) ring stage parasites. At ring stage, parasites were treated with E2G12 (700 ng/µl) under GIA condition for 24h. On the other hand, for rescue experiment, E2G12 was washed out and re-cultured in antibody free medium for 12h. Untreated control parasites, E2G12 treated parasites were harvested by washing twice with 1x PBS and thereafter fixing with 4% Paraformaldehyde (PFA) and 0.0075% Glutaraldehyde. Similarly, after rescue, parasites were harvested and fixed. After fixation, parasites were washed twice with 1x PBS and stained with DAPI (0.1µg/ml) for 15 min at RT. After nucleus staining, parasites were washed twice with 1x PBS and used for flow cytometry. Up to 0.5 million cells were assessed for the using LSR Fortessa (BD Biosciences, USA) and analyzed by FLOWJO software.

Untreated, Cerulenin and C75 treated HEK 239T and HCT 116 cultured cells were harvested after trypsinization and centrifugation. Cells were washed twice with ice cold 1x PBS and fixed with a fixing solution containing 4% Paraformaldehyde and 0.1% Glutaraldehyde for 2h. After fixation, cells were washed with 1x PBS and nucleus was stained with DAPI (0.1 µg/ml) for 15 min at RT. Cells were then washed twice with 1x PBS and then resuspended in equal volume of 1x PBS for flow cytometric analysis. Up to 80 thousand cells were assessed using LSR Fortessa (BD Biosciences, USA) and analyzed by FLOWJO software.

### Quantification and statistical analysis

Quantification and statistical calculation were done and plotted as mean ± S.E.M. Significance was calculated by unpaired Student’s t test or one-way ANOVA. Significance was considered as ∗*p* < 0.05, ∗∗*p* < 0.01, ∗∗∗*p* < 0.001, ∗∗∗∗*p* < 0.0001. Not significant was denoted as P=NS. N represents biological replicates. All quantitative experiments were repeated at least three times or more.

## Acknowledgements

We thank Prof. Sanjay A. Desai from NIAID, NIH, USA for providing pL6 and pUF1-Cas9 plasmids as a gift. We thank Prof. Christian Doerig from RMIT University, Melbourne, Australia, and Dr. Coralie Boulet and Prof. Mathieu Brochet from University of Geneva, Switzerland for providing transgenic PK6-HA and PK9-HA *P. falciparum* 3D7 parasite lines. We also thank Prof. Alain Verreault from University of Montreal, Canada, for initial discussion about parasite histone isolation and PTMs analysis. We are thankful to Dr. Sucheta Tripathi at CSIR-IICB for providing us Cyanobacteria strains. We thank Mr. Sayak Bhattacarjee, Mr. Satyaki Das and Mr. Syamantak Dey for their contribution in some figure preparations. We are thankful to MR4 BEI resources for Plasmodium parasite strains and other reagents. We thank Calcutta National Medical College, Kolkata, India for providing O^+^ human blood and human serum. We thank Eurofins Scientific, Bangalore, India for Transcriptomics services. We also thank Bioklone Biotech Pvt. Ltd, Chennai, India for the synthesis of custom-made anti-PfP2 mAb E2G12 antibody. We also thank Central Instrumentation facility at CSIR-IICB for helping with instrumentations particularly for in house Mass spectrometry facility ORBITRAP/qTOF. For fundings, we are thankful to Ramalingaswami Fellowship (BT/RLF/Re-entry/40/2016), Department of Biotechnology (DBT), Govt. of India, to SD; Core Research Grant (CRG/2018/000866), SERB, Department of Science and Technology (DST), Govt. of India to SD; Medical Biotechnology Research grant (BT/PR44703/BRB/10/2013/2021) Department of Biotechnology (DBT), Govt. of India, to SD and CSIR-IICB Institutional support through P07 and P50 grants. Funders had no intellectual inputs and did not perform any experiments.

## Authors Contributions

S.D.: Conceptualization, designing experiments, performed experiments (parasite transfection, parasite culture, biochemical and biophysical experiments), data analysis, figure preparation, manuscript writing and editing.

## NCBI data deposition

*P. falciparum* 3D7 and NF54 LncRNA nucleotide sequences are being deposited to NCBI database. Accession number will be provided when it is available.

## Lead Contact

For reagents and transgenic parasites, the lead contact

