## Supplementary figures and images for "Histone’s quality control by prokaryotic ClpP/ClpR regulates the eukaryotic mitotic cell cycle in malaria parasites"

### Supplementary Figure 1

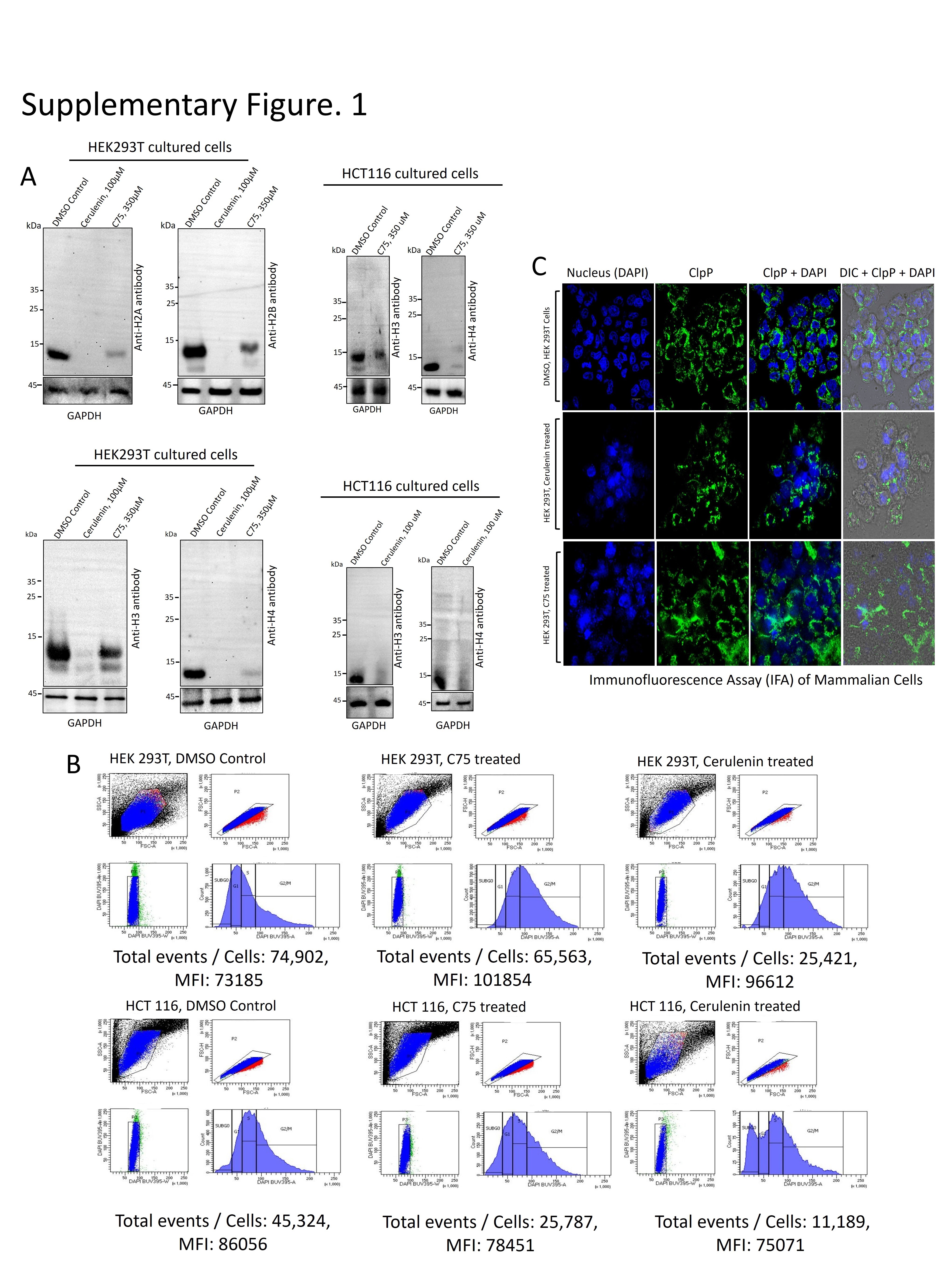

### Supplementary Figure 2

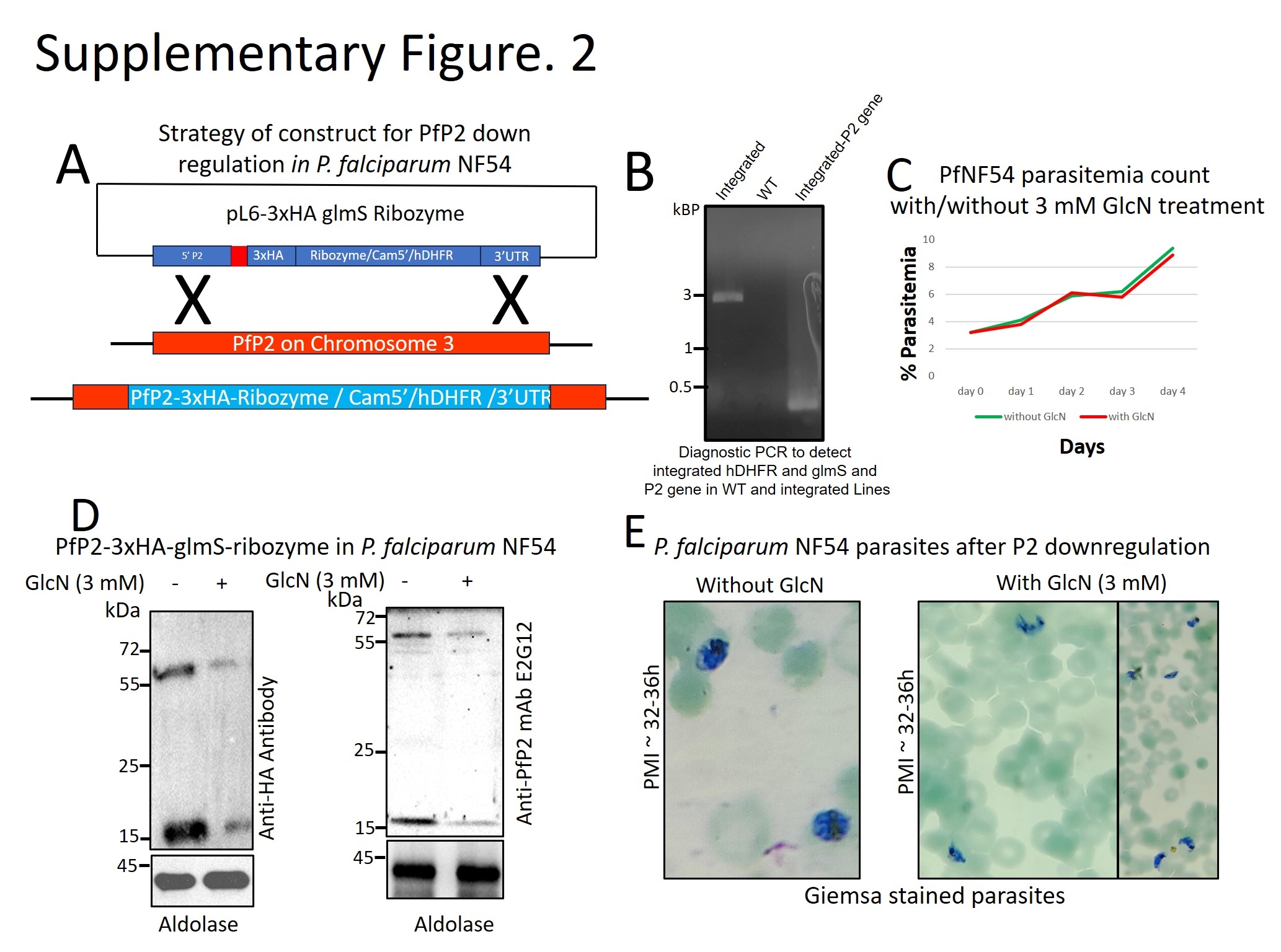

### Supplementary Figure 3

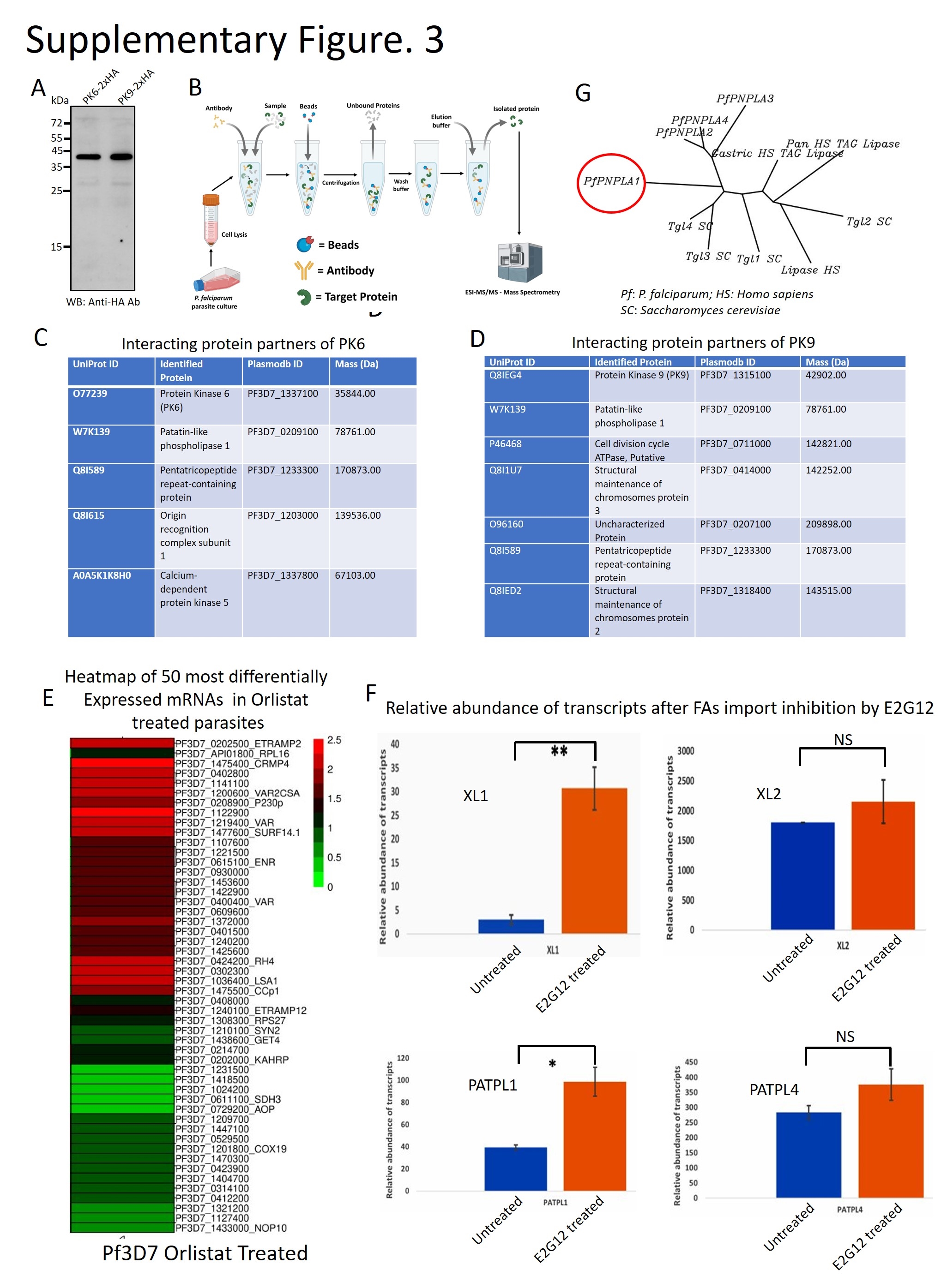

### Supplementary Figure 4

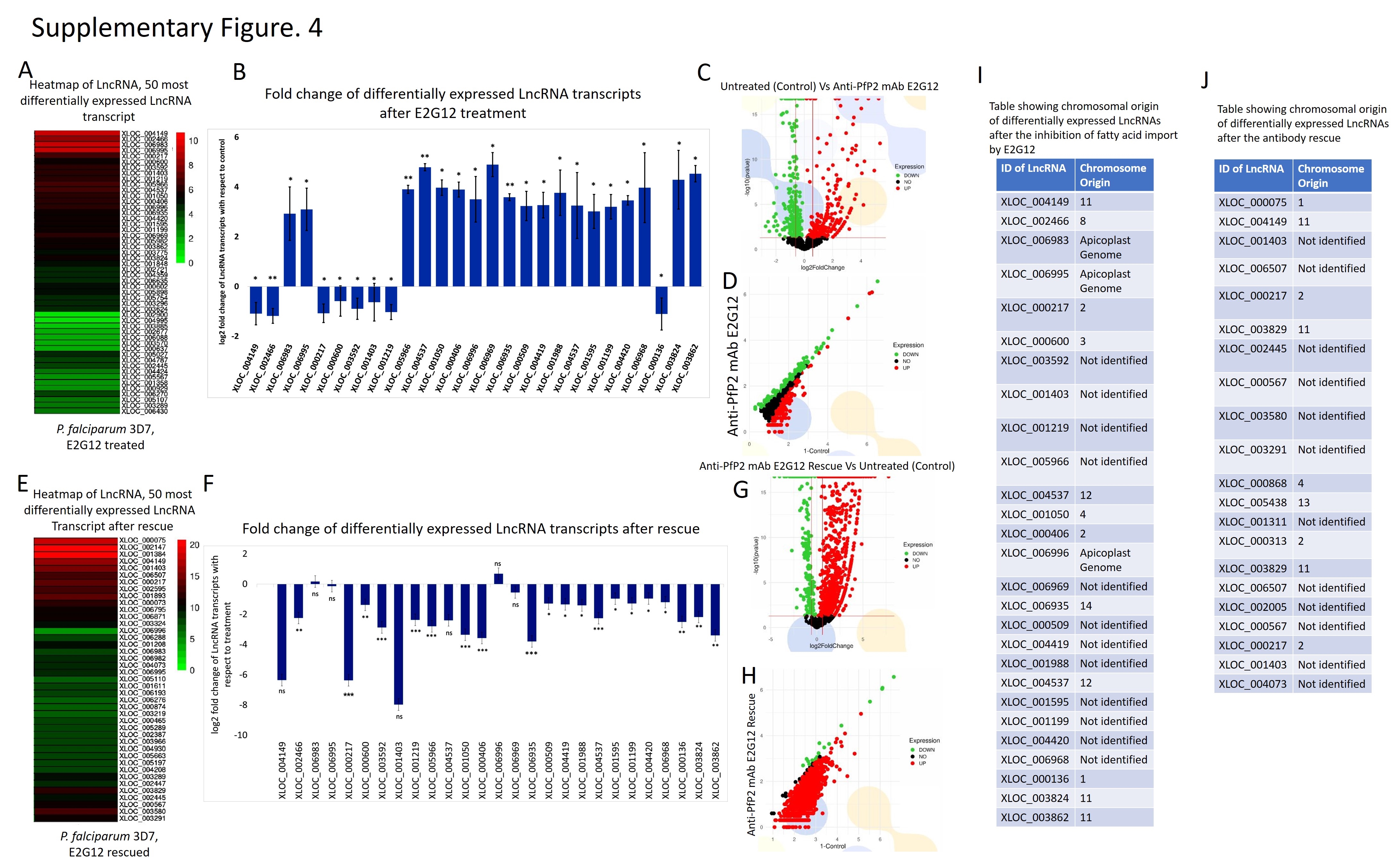

### Supplementary Figure 5

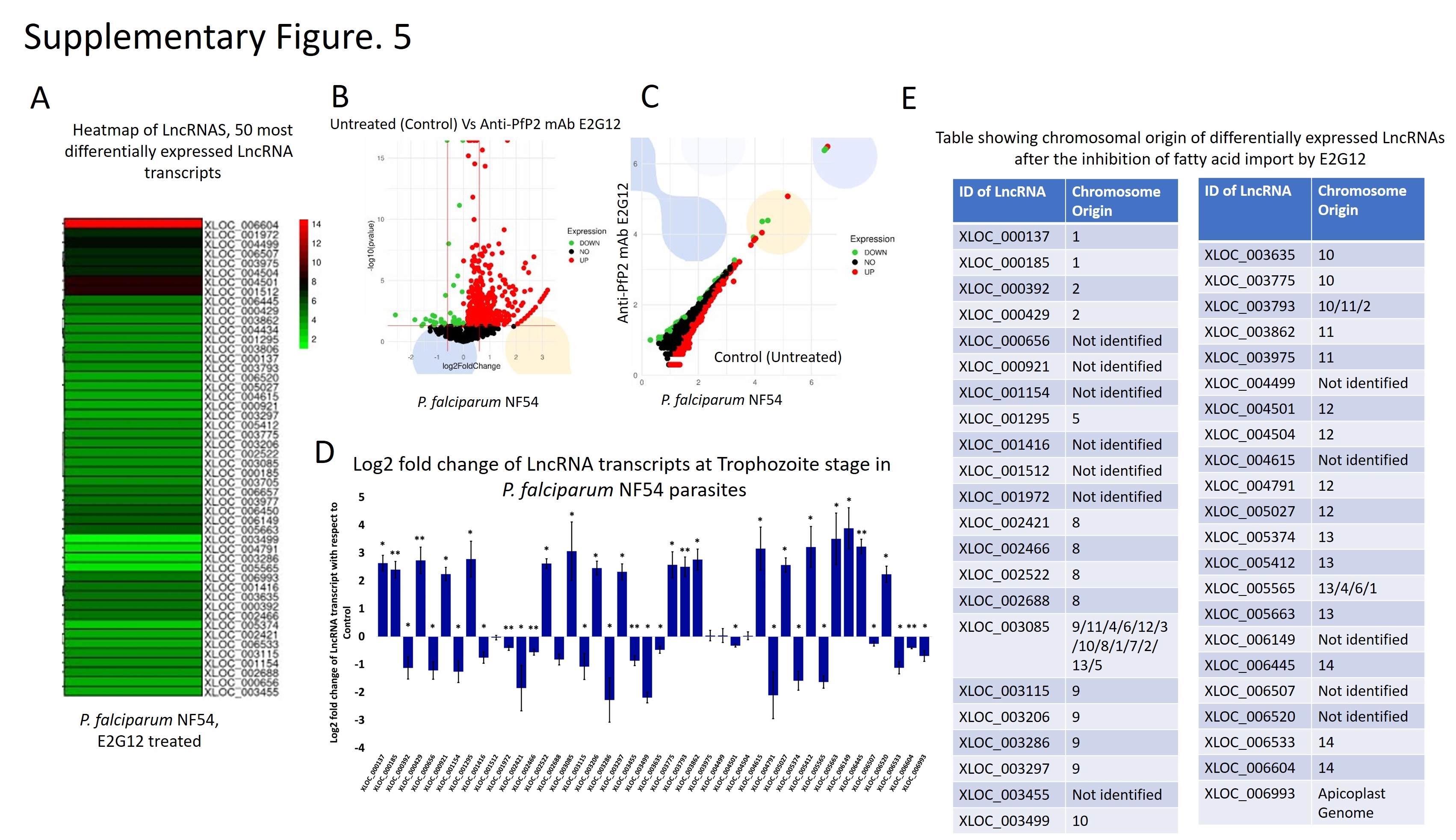

### Supplementary Figure 6

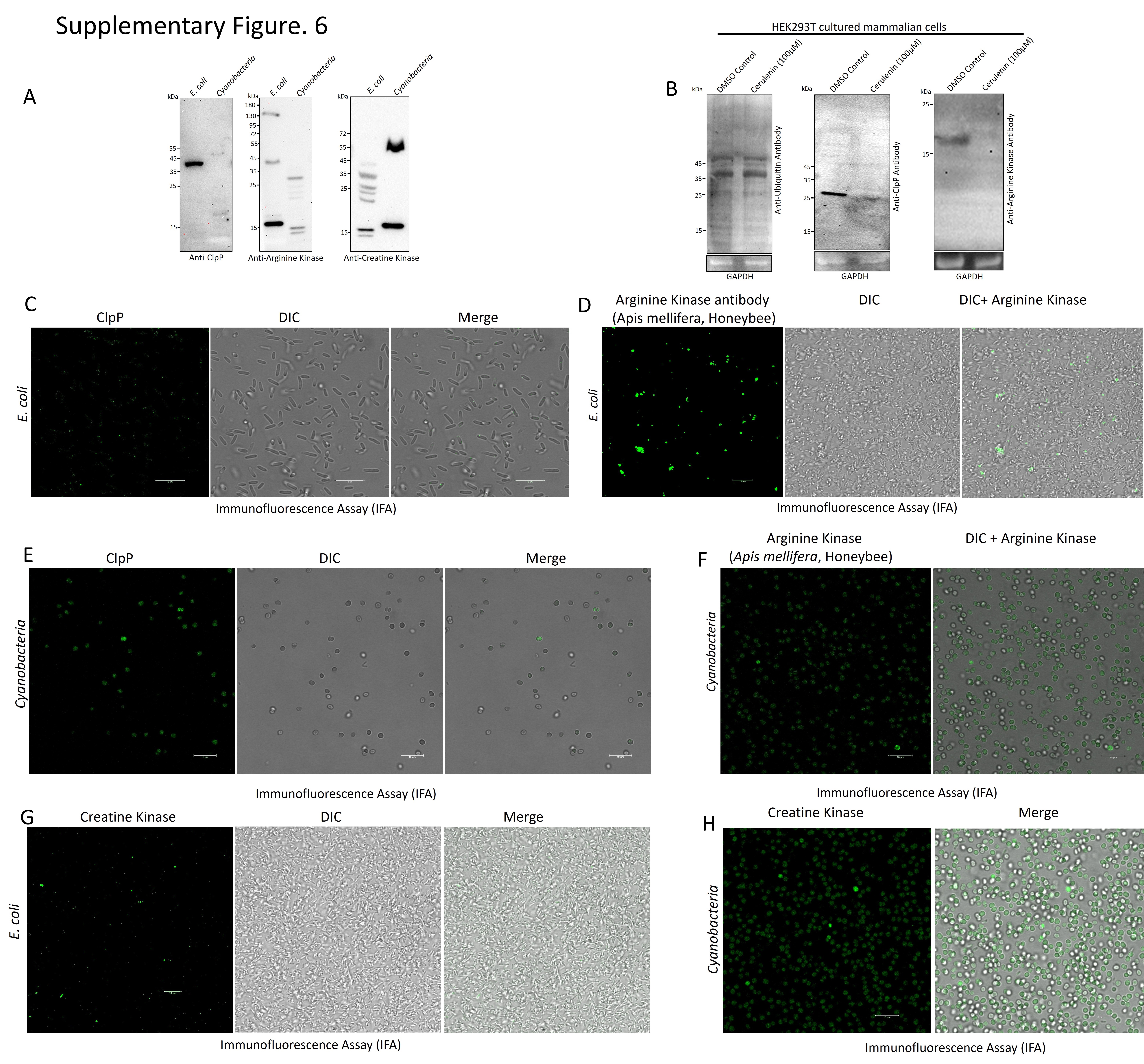

### Supplementary Figure 7

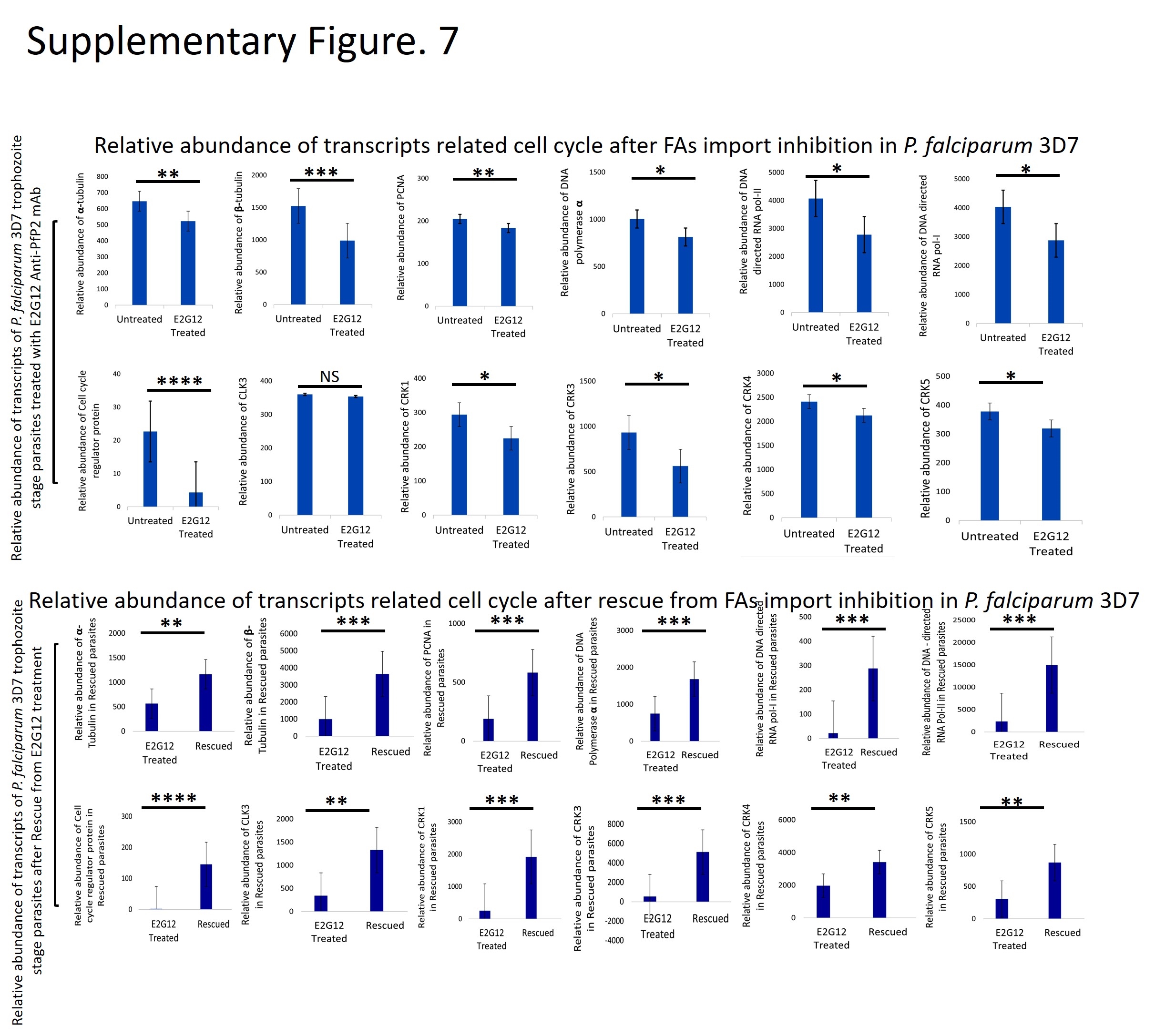

### Supplementary Figure 8

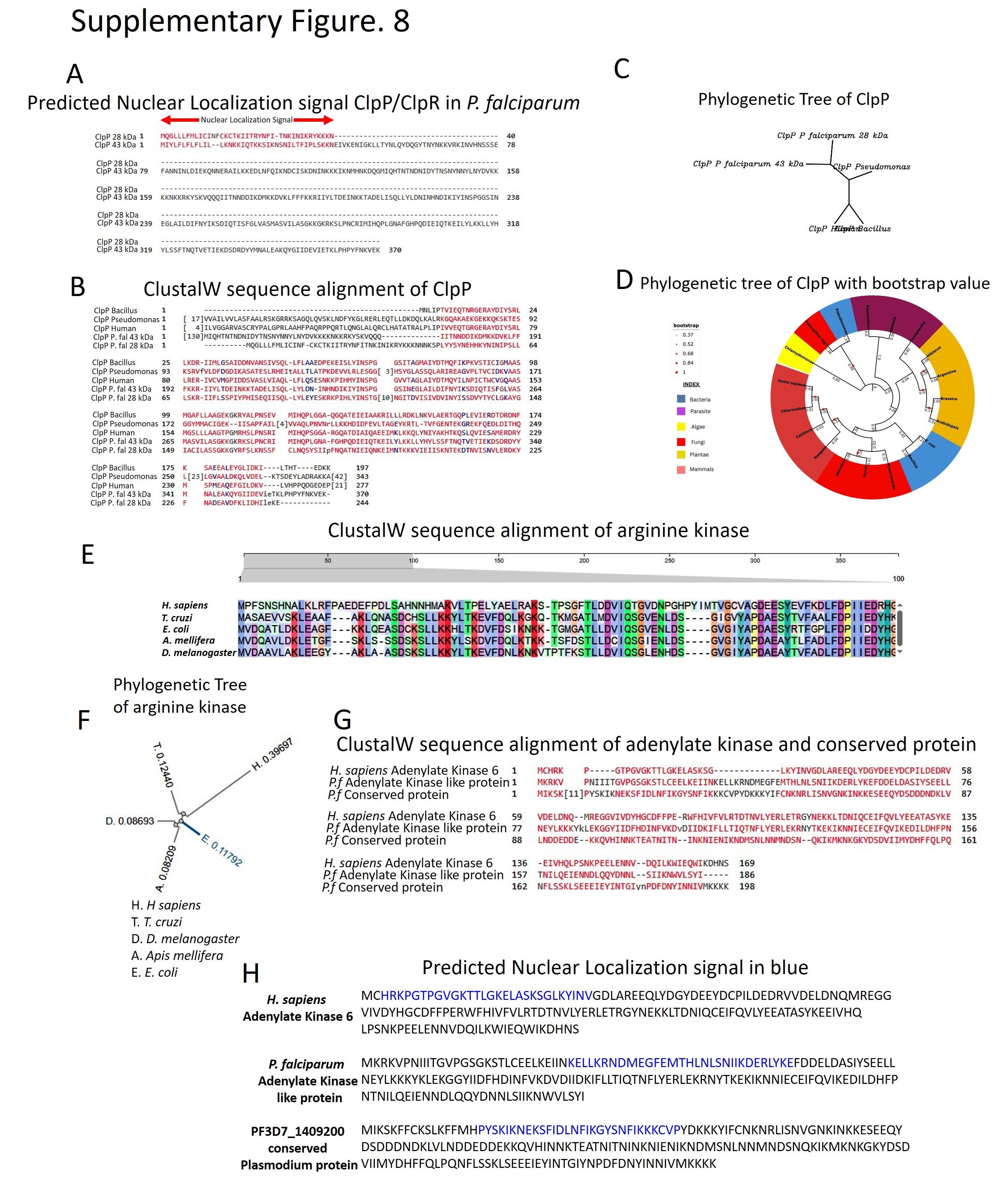
